# Limitations of *ad hoc* genotyping in detecting ash dieback tolerance in *Fraxinus excelsior*

**DOI:** 10.1101/2024.10.11.617393

**Authors:** Aglaia Szukala, Gregor M. Unger, Carlos Trujillo-Moya, Thomas Geburek, Thomas Kirisits, Silvio Schueler, Heino Konrad

**Author notes:** Corresponding authors: Aglaia Szukala.

## Abstract

Ash dieback (ADB), caused by the invasive alien fungal pathogen *Hymenoscyphus fraxineus*, is an emerging disease that poses major threats to the survival of common ash (*Fraxinus excelsior*) in Europe. Here, we report on a field trial aiming at the identification of ADB tolerant genotypes while encompassing the genetic diversity of common ash in Austria. Over 35,000 progenies from more than 600 putative tolerant mother trees were assessed for ADB symptoms over a period of three years. One tolerant and one susceptible progeny from a subset of the mother tree range (570 trees) were genotyped using the 4TREE array, a genotyping tool developed specifically to screen common ash provenances for their tolerance to ash dieback. We aimed to (1) better characterize the genetic structure of common ash in Austria, (2) identify hybrids between common ash and *F. angustifolia* (narrow-leaved ash), (3) detect genetic loci associated with tolerance to ADB, and (4) identify genetic variation associated with abiotic factors such as temperature and precipitation influencing the severity of ADB. We identify ash individuals from Eastern Austria matching putative hybrids with *F. angustifolia* showing varying degrees of *F. angustifolia* ancestry in structure analyses. The non-hybrid gene pool is characterized by shallow genetic structure following a west to east geographic pattern. In contrast to our expectations, genotype-damage association analyses overall fail to detect SNPs associated with tolerance to ADB. Our findings underscore that the polygenic architecture governing tolerance and the rapid decay of genomic linkage disequilibrium (LD) (within approximately 10 kb) significantly limit the ability to identify association outliers using the 4TREE SNP array. This highlights the constrained utility of chip genotyping approaches for this specific purpose. In contrast, the array proves successful in detecting hybrids between common ash and narrow-leaved ash, as well as outlier SNPs associated with abiotic predictors (such as annual precipitation and temperature) known to influence the severity of ADB damage.

## Introduction

Common ash (*Fraxinus excelsior*) is a widespread, ecologically important and one of the most valuable native European broad-leaved tree species for forestry. Due to its ability to thrive in diverse environmental conditions and particularly its considerable tolerance to drought (Boshier and Stewart, 2005; Dobrowolska et al., 2011), common ash has been regarded to cope well with increasing warm-dry conditions forecasted in the future due to climate change (Dyderski et al., 2018). However, since the early 1990s a significant decline of common ash in Europe has been caused by the ascomycete fungus *Hymenoscyphus fraxineus* (synonym *Chalara fraxinea*), leading to an emerging disease known as ash dieback (ADB) (Gross et al., 2014; McKinney et al., 2014; McMullan et al., 2018; Marçais et al., 2022). The pathogen is native to East Asia and has been spreading since its first observation in 1992 in Poland into the majority of the common ash distribution range, and it is now known to occur in 32 European countries (Marçais et al., 2022).

Since the emergence of ADB in Europe, the populations of *F. excelsior* have been experiencing high and unprecedented mortality rates (Marçais et al., 2017; Coker et al., 2019; Madsen et al., 2021; Matisone et al., 2021; George et al., 2022). If the loss of common ash in deciduous forests proceeds at the current rate, this would lead to a lasting change in forest ecosystems and potentially drive organisms tightly associated with ash to extinction (Mitchell et al., 2014; Coker et al., 2019; Díaz-Yáñez et al., 2020; Marçais et al., 2022). Despite research suggesting that climate change may partly reduce the loss of common ash through ADB due to an increase of areas with less favorable environmental conditions for *H. fraxineus* (i. e. a warm/hot-dry climate) (Goberville et al., 2016; Marçais et al., 2022), studies underpinned the necessity for fast action towards the rescue of common ash in Europe (Pautasso et al., 2013; George et al., 2022).

A rapidly increasing body of research monitored the development and advance of ADB in Europe since its appearance (Vasaitis and Enderle, 2017), detecting sporadic (1%–5%) but genetically determined and heritable high tolerance to ADB in many European countries (Kirisits and Freinschlag, 2012; Kjær et al., 2012; McKinney et al., 2014; Lobo et al., 2015; Landolt et al., 2016; Pliūra et al., 2017; Marçais et al., 2022). The terms tolerance and resistance have been used to characterize the variation of damage caused by ADB on common ash individuals. Following Landolt et al. (2016) and Marçais et al., (2022, 2023), tolerance is used in the present study, as *H. fraxineus* can fulfill its life cycle and thus form fruiting bodies on leaves of ash trees with low or high damage levels on woody parts (on which sporulation of the pathogen occurs only rarely), which agrees well with the concept of disease tolerance in plant pathology.

Studies attempted to better understand the genetic basis of ADB tolerance in Denmark (Harper et al., 2016), Germany (Enderle et al., 2017), Poland (Meger et al., 2024), Sweden (Chaudhary et al., 2020), the UK (Stocks et al., 2019; Sollars et al., 2017), and across Europe (Doonan et al., 2023). Associative transcriptomics (Harper et al., 2016) found expressed loci associated with tolerant *versus* non-tolerant phenotypes. A screening of whole-genome methylation variation identified over 1,600 genomic regions differentially methylated between susceptible and tolerant individuals, with a possible role in dosage regulation of susceptibility related genes (Sollars and Buggs, 2018). Moreover, Stocks et al. (2019) identified over 3,000 SNPs associated with ADB damage using pooled whole-genome sequencing, enabling prediction of ADB presence/absence with high accuracy.

Following these achievements and recent advancements in the assembly and annotation of whole ash genomes (Sollars et al., 2017; Huff et al., 2022), the 4TREE SNPs array was developed, incorporating informative SNPs for ADB tolerance, along with putatively neutral loci (Guilbaud et al., 2020, *in prep.*). This resulted in the creation of a cost-efficient, high-throughput genotyping tool aiming at facilitating collaboration among breeding programs across Europe. Specifically, SNPs were chosen based on their strong association with ash dieback tolerance, as determined in prior studies (Harper et al., 2016; Sollars et al., 2017; Stocks et al., 2019). With a relatively small genome size (approximately 880 Mbp, Sollars et al., 2017), the 4TREE array holds promise as a valuable resource for characterizing informative genetic markers related to ADB tolerance in common ash.

The present study aimed to leverage these recent advancements to identify genotypes with heightened tolerance to ADB through a screening of a comprehensive and diverse assemblage of common ash specimens collected across Austria in the course of the breeding program “Ash in distress” (Unger et al., 2021; Fig. 1A). We also aimed to detect hybrids between common ash and *F. angustifolia* (narrow-leaved ash), as hybridization between these two species is common in areas where their distribution ranges overlap (Fernandez-Manjarres et al., 2006) and has been reported to occur in Austria, particularly in forests along the eastern stretch of the Danube and other rivers in Eastern Austria (Zukrigl, 1997; Heinze et al. 2017). However, identifying hybrids or introgressed individuals in natural populations is challenging, since the range of morphological variation of the hybrids largely overlaps with the parental species (Thomasset et al., 2011). Notably, misclassifying introgressed individuals can impede the accurate identification of genetic loci associated with tolerance in association analyses. Finally, we aimed at an overall characterization of the distribution of potentially adaptive genetic variation of common ash in Austria focusing on key abiotic factors like altitude, precipitation and temperature. This exploration aims to deepen our understanding of the complex interplay between environmental conditions and genetic traits, shedding light on the mechanisms underlying tolerance to ADB.

**Figure 1.**
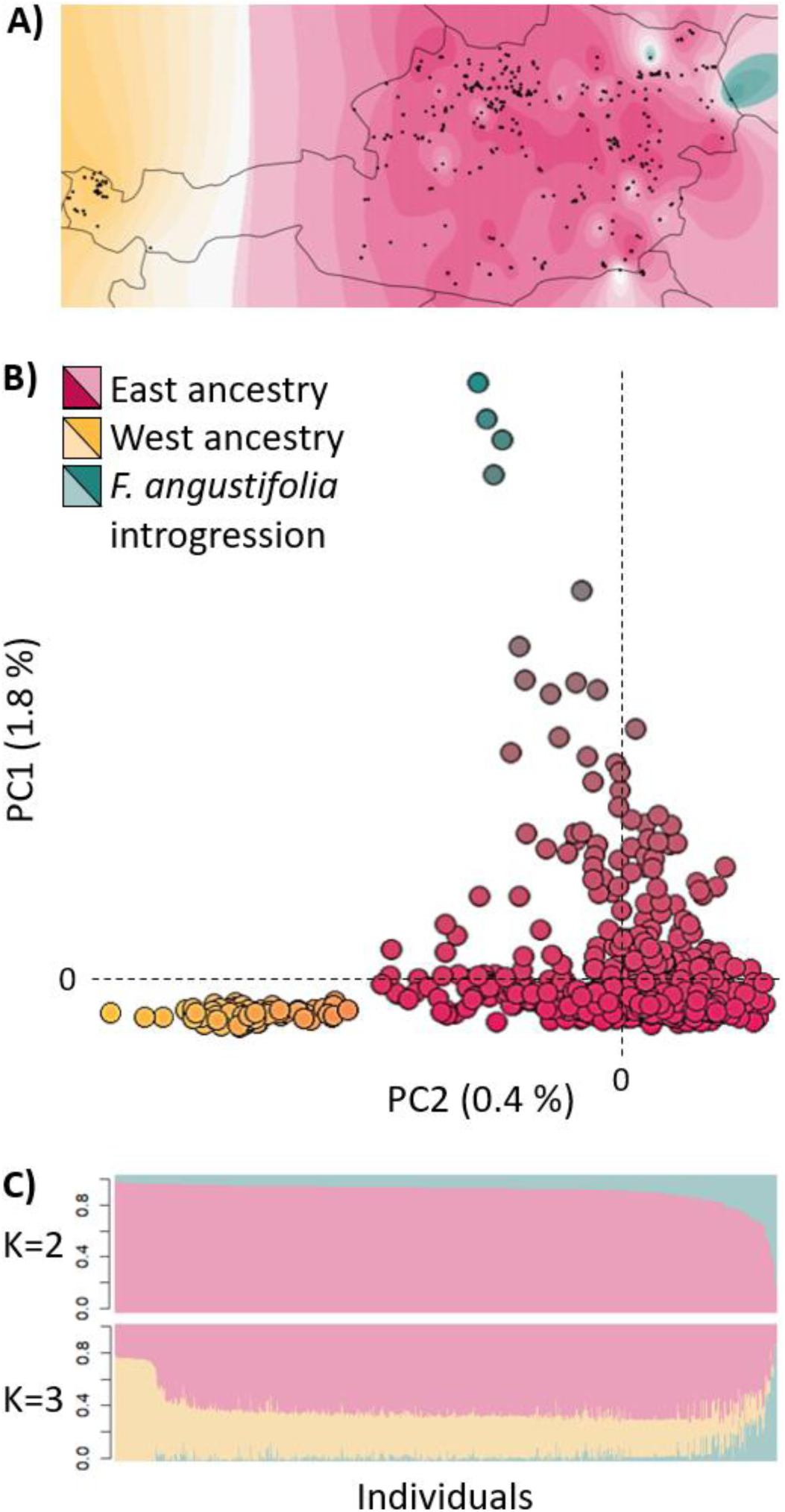
Map of Austria showing the sampling locations of 570 ADB tolerant trees collected (black dots), the progenies of which were analyzed in this study. The color shades of the map correspond to the clusters detected in the PCA and admixture analyses. Note that in the easternmost part of the country the green to white shades denote individuals with different proportions of *F. angustifolia* introgression **(A)**. Principal component analysis showing samples with *F. angustifolia* introgression along the first component (PC1) and a west to east genetic diversity gradient along the second component (PC2) **(B)**. Admixture analysis performed using TESS3 (best K = 3) similarly shows *F. angustifolia* introgression, as well as an east-west cline of genetic diversity **(C)**.

## Material and Methods

### Plant material and experimental design

To establish a sufficient breeding base of resistant clones without diminishing the gene pool, we selected 611 putatively ADB tolerant and seed bearing single trees (i. e., functionally females; Saumitou-Laprade et al., 2018) of *F. excelsior* distributed in forests across Austria. Forest stands were selected based on the rare presence of remarkably healthy or only slightly damaged trees despite these being surrounded by a high number of infected and heavily damaged or dead individuals, whereas sites in rural settings or in areas with low disease incidence were avoided because of too low infection pressure. Likewise, 25 seed bearing ash clones were selected in the clonal seed orchard in Feldkirchen an der Donau (Upper Austria) and 46 in the clonal seed orchard in Bad Gleichenberg (Styria) (Kirisits and Freinschlag, 2012; Freinschlag, 2013). ADB damage levels varied greatly among genotypes in the two seed orchards. Moreover, one tolerant genotype from a clonal seed orchard in Denmark and two *Fraxinus pennsylvanica* trees in forests in Austria were chosen. However, only common ash trees from Austria were considered for genotyping in the present study.

Seeds were harvested in 2015 and 2017 from the selected ash trees (the majority of which being considered as putatively ADB tolerant), and 35,718 seedlings germinated and were reared in plastic multi-container blocks in the experimental nursery of the Austrian Research Centre for Forests (BFW) in Tulln an der Donau (Lower Austria). In order to systematically assess the susceptibility to ADB over a three-year span, the one-year-old offspring seedlings (on average about 50 seedlings per mother tree) were arranged in a field trial in a completely randomized block-design. Seedlings were arranged in rows, with distances of 0.8 m between rows and 0.4 m between seedlings within rows. To increase the level of natural infection likelihood and thus the selection pressure on the seedlings, ash leaf litter (i. e., pseudosclerotial ash leaf material collected in heavily infested ash stands) was placed between seedlings in the respective years of the installation of the various parts of field test.

*Fraxinus excelsior* is characterized by complex seed dormancy; thus seeds harvested in 2015 germinated during the two following years, in 2016 and 2017, while those harvested in 2017 germinated in 2018 and 2019. Thus, the field trial included four sets of seedlings germinated during four subsequent years (2016 to 2019) and planted in four separate, but adjacently arranged sub-trials, named hereafter R1 (established in 2017), R2 (established in 2018), R3 (established in 2019) and R4 (established in 2020; not investigated in the present study). Progeny of field tests R1, R2, R3 and R4 were assessed for their susceptibility to ash dieback (intensity/degree of ADB) annually over a period of three years (results for R4 are not shown here). Leaf samples were collected from all offspring seedlings during the year of field trial installation, dried on silica gel and stored at room temperature until DNA extraction. For the present study, offspring from 570 ash mother trees in the sub-trials R1, R2 and R3 were selected for genotyping (Fig. 1A).

### Phenotyping

For the damage assessment on young ash plants only woody parts were visually inspected for symptoms of ADB and rated in six damage classes (DC; DC 1, 0 % ash dieback, not diseased; DC 2, >0 to 10%; DC 3, >10 % to 50 %; DC 4, >50 % to 90 %; DC 5, >90 % to <100 % and DC 6, 100 % damaged, dead; Fig. S1A; Unger et al., 2021). The ADB intensity is thus expressed as the percentage of damaged woody parts in relation to the tree height. Care was taken to distinguish damage due to ash dieback from other causes of damage as best as possible, e. g. establishment failure in the year of planting. We kept track of the number of individuals belonging to each damage class on a yearly basis. Where possible, one completely healthy progeny (i. e. DC = 1) and a dead one (i. e. DC = 6) from each of the 570 mother trees were chosen for genotyping at the end of the third, second, and first assessment period in the R1, R2 and R3 field trials, respectively. Where no healthy or dead individuals were available, individuals of intermediate damage classes were chosen instead. For identifying diseased genotypes, the final scoring of ADB intensity (third year of documentation) was used for all genotyped samples.

We also collected data regarding flushing behavior of the progenies to distinguish early from late flushing individuals (Table S1). In the year after planting (i. e. in 2018 in R1, in 2019 in R2 and in 2020 in R3), each seedling was scored by applying a slightly modified scheme developed by Andrić et al. (2016) for *F. angustifolia* from dormant buds (Flush 1) to completely unfolded, smooth and shiny leaves (Flush 6). Leaf flushing classification started in early April and was repeated after a period of two weeks. The association between the categorical variables “field trials” (i. e. R1-R3) and first or second Flush with DC was tested in R v. 4.2.2 using Pearson’s Chi-squared test and visually inspected by means of Cohen-friendly association plots for categorical variables of the vcd package v. 1.4-12 (Friendly and Meyer, 2015).

### DNA extraction and genotyping

DNA of the selected progenies was extracted at the Austrian Institute of Technology (Tulln an der Donau, Lower Austria) using the Qiagen DNeasy 96 Plant Kit according to the manufacturer’s recommendation. DNA quality and quantity were measured using PicoGreen™ dsDNA Assay Kits (ThermoFisher). We conducted genotyping of a total of 1,107 individuals, encompassing both symptomless and diseased progeny from the majority of initially selected mothers, at Thermo Fisher Scientific Inc. using the 4TREE Axiom SNP array (Guilbaud et al., *in prep*), which encompasses a set of 13,407 SNPs. The design of the SNP array was based on the *F. excelsior* genome assembly BATG-0.5 (BioProject number PRJEB4958, Sollars et al., 2017). In light of the later publication of a new chromosome-level assembly (FRAX_001 genome, BioProject number PRJNA713541, Dong et al., 2022), probes of the 4TREE array were aligned to the updated reference genome using bowtie v. 2.3.5.1 (Langmead and Salzberg, 2012) to retrieve SNPs positions in the new reference genome, enabling e.g. more accurate estimates of linkage disequilibrium (LD).

### SNPs filtering

The raw data was filtered to retain high quality SNP calls using Axiom Analysis Suite v. 5.2.0.65 (Thermo Fisher Scientific Inc.) using the cut-off options recommended by Thermofisher as reported in Table S2. Previous to association analyses we further excluded monomorphic loci using the *-c 1* option in bcftools v. 1.18 (Danecek et al., 2021), removed rare variants using a minimum 0.05 minor allele frequency (MAF) cutoff, and allowed for maximum missing rate 0.01 using vcftools v. 0.1.16 (Danecek et al., 2011) on the vcf file of the quality-trimmed SNPs.

### Genetic structure analyses

We inferred spatial population structure of the genotype data using principal component analysis (PCA) and a spatial genetic structure method implemented in TESS3 (Caye et al., 2016) using the full dataset and randomly retaining one offspring per mother. These analyses were performed in R using the packages vegan v.2.6-4 (Oksanen et al., 2012) and tess3r v. 1.1.0 for PCA and admixture analyses, respectively. TESS3 was run for values of K from 1 to 10 and the K best explaining the variance observed in the data was evaluated visually using the cross-validation score, as well as by comparing these values with the total amount of variance explained by the first 10 principal components in PCA.

### Analysis of linkage disequilibrium decay

We estimated LD using TASSEL v. 20230110 (Bradbury et al., 2007) by calculating the squared allele frequency correlation coefficient (r^2^) between all pairs of markers (*Full Matrix* option) in each chromosome separately and setting heterozygous calls to missing. We then visualized the resulting LD estimates and computed distance at which LD falls to half of its maximum value using the qqman v. 0.1.9 CRAN package in R (Turner, 2018). Lastly, we utilized the Bioconductor chromPlot package (Verdugo and Orostica, 2017) to plot the by-chromosome estimated LD decay length up- and downstream of each SNP position to visualize and estimate the total amount of chromosomal regions that may be informative on genotype-trait associations.

### GWAS of ADB and flushing behavior

We conducted GWAS on DC, which was coded both as a binary and ordinal phenotype. For the binary analysis, we retained only the most extreme phenotypes: DC 1 (fully healthy) and DC 6 (dead). In an alternative approach, classes 1 and 2 were pooled as healthy, while classes 5 and 6 were pooled together and considered susceptible. Additionally, DC was treated as an ordinal categorical variable with six levels (1 to 6, as described previously). Missing genotypes were imputed using the most common genotype at each locus across individuals.

For the binary phenotype, GWAS was performed using logistic regression implemented in the CRAN R package glmnet (Friedman, Hastie, and Tibshirani, 2010). Initially, a naive association test was conducted using Fisher’s exact test without correcting for population stratification. Subsequently, a lasso binomial logistic regression model was fitted using the *cv.glmnet* function, specifying *family=”binomial”*, *alpha=1*, and *lambda.min.ratio=0.01.* The SNP matrix was corrected for population stratification by testing K values ranging from 2 to 7. When DC was coded as an ordinal categorical variable, analysis was conducted in glmnet using the function *cv.glmnet* and specifying *family=”multinomial”* and *alpha=”1”* to implement a lasso regression. After identifying the best lambda value through cross-validation, a final model was fitted using this optimal lambda via the *glmnet* function.

For all glmnet models, SNPs with non-zero coefficients at the best lambda were considered potentially associated. These SNPs were validated through class prediction using the *predict* function, specifying *type=”response”* or *type=“class”* to generate predicted probabilities or classes, respectively. Prediction accuracy was assessed using a confusion matrix, calculated as the proportion of correctly classified observations relative to the total number of observations. Additionally, we performed discriminant analysis of principal components (DAPC)-based feature selection using the SNPs with non-zero coefficients to test how well these can discriminate between DC categories.

In cases where DC was treated as a quantitative response, we employed both generalized linear models (GLM) and mixed linear models (MLM) to account for confounding effects of genetic structure using TASSEL. The GLM utilized the first two principal components (PCs) from a PCA analysis to correct for population stratification (note: varying the number of PCs from 2 to 10 did not affect the results). We then ran the GLM using default settings with 100,000 permutations. For MLM analysis, relatedness between individuals was estimated using a kinship matrix, generated via the *centered_IBS* method (Endelman and Jannink, 2012) to account for random effects by incorporating kinship into the model. Additionally, we also run a model including the field trials design (R) as an additional random effect. All the listed approaches were also applied to analyze the association between genotype and the first and second flushing using multinomial logistic regression and linear models. A Bonferroni correction was applied to the p-values, and a significance threshold of 0.05 was used to detect significant outliers.

### Genotype-environment associations (GEA)

We tested the association between genotypes and environmental variables using latent factor mixed models (lfmm) in the R package lfmm2 v. 1.0 available on github (Caye et al., 2019). This method models population structure and other hidden confounding factors as an unobserved (latent) variable (Frichot et al., 2013; Caye et al., 2019). Missing genotypes were imputed using the most common genotype at each locus across individuals. Previous to the analysis, quantitative predictors were tested for collinearity using Pearson’s correlation and visual inspection. We used the regularized least squares estimation method (function *lfmm_ridge*) with K values ranging from 2 to 7 and the genomic inflation factor (λ) to adjust the significance of p-values. The effect of the estimated lambda on the p-values distribution was checked visually by means of Q-Q plots. We tested several models incrementally including environmental predictors reported in Table S1 (Supplementary Information). Climatic data was obtained from the WorldClim database (Fick and Hijmans 2017) using the geographic coordinates of the sampled mother trees and the *worldclim_tile* function from the geodata package in R (Hijmans et al. 2023). We chose to analyze a set of climatic variables that were previously reported to have an influence on the intensity of ABD (Table S1), such as precipitation (Chumanová et al., 2019; Pliūra et al., 2015; Marçais et al., 2022). Additionally, we ran PCA of the environmental variables and used the first three synthetic PC axes of the PCA as predictors instead of the single environmental predictors (i. e. genotype ∼ PCs of environmental predictors + latent factors + E). Significant outlier SNPs were defined as having a false discovery rate (FDR) < 0.05.

Further, we used the multivariate method redundancy analysis (RDA) (Dray et al., 2012; Bourret et al., 2014; Forester et al., 2018) implemented in the R package vegan to further explore the association between the genotype and the environment. In contrast to univariate tests which need FDR correction, RDA treats genetic markers as a multivariate response, allowing for easier discovery of weak, multilocus molecular signatures of selection (Capblancq et al., n.d.; Rellstab et al., 2015). Missing genotypes were imputed in the same way as in the gwas and lfmm analysis. We used the first 2 to 7 PC axes of the genotype matrix as a proxy for population structure and the first synthetic PC axis of the PCA of environmental variables by implementing the formula genotype ∼ PC of environmental predictors + condition (PCs). The significance of the model was tested using an ANOVA like permutation test (Legendre et al., 2011) with 999 permutations using the function *anova.cca()* and setting the option *by=”axis”* to test the significance of each constrained RDA axes (Legendre et al., 2011). RDA scores measuring the association between the genotype and the predictors were assigned to each SNP and candidate SNPs were defined as being above the significance threshold ± 3 standard deviations (SD) from the mean score of the constrained axis. The potential role of the genes containing outlier SNPs in the adaptation to the environment was interpreted based on the known gene function.

## Results

### ADB damage assessments

ADB intensity increased in each of the sub-tests during the respective three-year-long assessment periods (Fig. S1B). While 82 % (4,973 of 6,033 active plants) of progenies in R1 were still undamaged (DC 1) by ADB at the first assessment, this proportion decreased to 58% in the second assessment year and was only 24 % (corresponding to 1,449 of 6,032 active plants) in the third year. The R2 and R3 test sites included the highest and lowest number of progenies, respectively, with 23,433 and 3,785 progeny plants, respectively, at the beginning of the assessments. In R2 and R3 the proportion of progenies of DC 1 changed from 55 % and 81 % in the first assessment year to 22 % and 31 % in the third assessment year, respectively. Notably, in R3 the proportion of progenies in DC 1 remained stable in the second and third years of assessment.

### Association between germination year, flushing behavior and the degree of damage

We observed that the categorical variables R (i. e. the field test sites R1-R3) and Flush (both first and second rating) were associated with DC (Pearson’s Chi-squared p < 2.2e-16, p = 6.033e-13, and p = 0.0003337, for R, first and second Flush, respectively, Fig. S2). We found an excess of individuals of DC = 1, 4, 5, and 6 in R1, and an excess of individuals of DC 2 and DC 3 in R2 and R3 (Fig. S2A). To minimize the effects of the test sites on the discovery of outlier SNPs in association analyses we performed additional rounds of association analyses, retaining only individuals from R2 (i. e. the largest test site) including the R variable as a random effect in MLM. The DC also showed a stronger correlation with the flushing behavior at the second assessment, where we observed an excess of individuals of DC 6 in the category Flush 1 (Fig. S2C). Thus, a high proportion of individuals that were dead by the end of the third assessment period were characterized by late flushing (i. e. being in the winter bud stage even at the second flushing assessment) in the year after planting. In contrast, symptomless individuals (DC 1) were more frequently early flushing.

### Filtering of genotypes reveals unexpectedly high amounts of monomorphic sites and rare alleles

We genotyped 1,107 progenies at 13,407 SNP loci included in the 4TREE array. After filtering SNPs based on the quality call thresholds recommended by Thermofisher 11,582 SNPs and 1,096 individuals were retained. A further SNPs filtering step excluded loci that were monomorphic, had a MAF < 0.05 and a missing rate of 0.01. After applying these thresholds, we retained 7,313 SNPs for 1,037 individuals (Table S1), meaning that ca. 37 % of the retained loci could not be included in downstream analyses, as being non-informative because of too low MAF (2,669 sites), or they were not supported by enough individuals (1,381) or monomorphic (219 sites). This trimmed dataset excluded a very high proportion of the 1,704 candidate SNPs included in the 4TREE array and previously identified by Stocks et al. (2019): We retained only 192 out of the 1,704 original outlier SNPs, as the majority of them was revealed to have too low MAF in our dataset.

Moreover, after blasting the 4TREE SNPs to the chromosome-level genome assembly, we found that out of 13,407 SNP only 9,910 SNPs had unique hits, while 2,429 SNPs were mapped to more than one position, and 1,068 did not map to any position in the new reference genome. We checked which of the 7,313 SNPs that passed filtering had a unique hit to the chromosome-level reference genome and retained 6,742 SNPs, including 179 (10.5 %) candidates from the Stocks et al. (2019) study, that were used for LD decay estimation and association analyses.

### Weak genetic structure and introgression with the closely related *F. angustifolia*

The PCA analysis (Fig. 2B) revealed weak genetic structure of common ash in Austria, as exemplified by the low amount of variance explained by the first two principal components (i. e. 1.8 % in PC1, and 0.4 % in PC2). Both admixture and PCA analyses identified K=3 as the number of clusters best fitting the data (Fig. S3A and B). Interestingly, the first principal component of the PCA consistently with K=2 of the admixture analysis (Fig. 2C) reveals some differentiated individuals sampled in Eastern Austria (Fig. 1A) that were previously identified as putative *F. angustifolia* or hybrids between common ash and *F. angustifolia* based on morphology (authors’ personal observation). More specifically, 59 individuals were found to have between 10 % and 90 % of *F. angustifolia* ancestry (Table S3A). We identified 75 SNPs showing the strongest separation between *F. excelsior* and *F. angustifolia* by means of RDA using the first PC as constraining variable and defining as “introgression outliers”, those loci being ± 3.5 SD from the mean score of the constrained axis (Table S3B). Finally, in PCA and admixture analyses, we consistently observed that genetic diversity follows a west to east geographic separation (PC2 and K=3).

**Figure 2.**
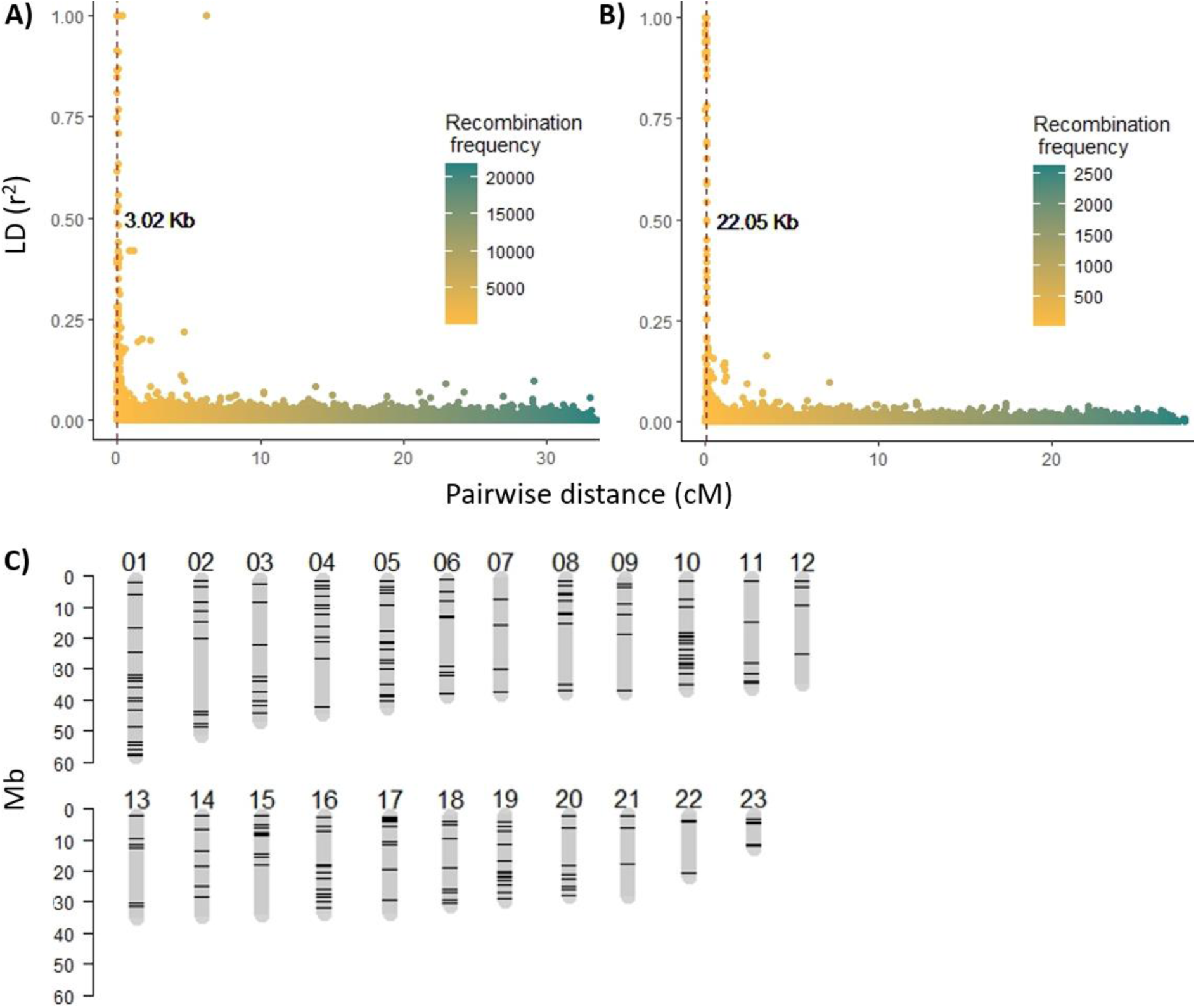
Rate of LD decay in chromosome 12 where the decay distance is shortest **(A)** and chromosome 19 where it is the largest **(B)**. The color scale shows the recombination frequency for a specific genomic distance expressed in centiMorgan (cM). The vertical dashed line and value reported in bold indicate the distance at which LD falls to half of its maximum value **(A, B)**. Graphical representation of the 23 chromosomes showing as black bands the genomic regions potentially covered in association analyses using the 4TREE array (after data quality trimming) as being within the range of by-chromosome estimates of LD decay **(C)**.

### Patterns of LD decay

After detection of hybrids between common ash and *F. angustifolia* we removed individuals having over 10 % *F. angustifolia* ancestry (Table S3) from the dataset for subsequent analyses. We found that the rate of LD decay is very fast and varies across chromosomes (Fig. 2 and S4), with average decay being at 9.7 Kb (Fig. S4). The decay distance is shortest in chromosome 12 (3.02 Kb, Fig. 2A), while largest in chromosome 19 (22.05 kb, Fig. 2B). Based on the estimates of by chromosome LD decay, the extent of genomic regions being in LD with SNPs after data pruning and thus informative in association analyses varied between 5.3 % of total chromosomal length in chromosome 12 to 30.1 % in chromosome 19 (Fig. 2C).

### Association analyses fail to detect candidate SNPs associated with DC

GWAS analyses were run after exclusion of individuals having over 10 % *F. angustifolia* ancestry. No significant associated SNPs were detected for the predictor DC using binomial logistic regression in glmnet. Using multinomial logistic regression, we identified 13 putatively associated SNPs having non-zero coefficients (including one SNP that was previously identified by Stocks et al., 2019), but validation of these SNPs by means of phenotype prediction based on the same model and DAPC cross-validation failed (prediction accuracy = 0.012, Fig. 5, 6).

Similarly, we were unable to detect significant SNPs associated with DC using the GLM (Fig. S7A,B) and the MLM model (Fig. 3A, S7C,D) in TASSEL. The addition of kinship and R as random effects did not significantly improve the model (Fig. 3A). Moreover, we observed a deviation from the expected distribution of p-values in the Q-Q plot of the raw p-values indicative of an excess of false negatives (Fig. S7A,D). This bias was not corrected by the inclusion of more PCs as fixed effect (genetic structure) or kinship and R as random effects. Although none of the SNPs had a p-value above the set Bonferroni correction threshold, both GLM and MLM models detected two SNPs with low p-value (Table S4) on chromosome 3 and 4 (Fig. 3A, S7B-D). Although these loci, along with those identified by multinomial logistic regression in glmnet, may contribute to immune defense responses, they clearly account for only an infinitesimal portion of the complexity underlying tolerance to ash dieback.

**Figure 3.**
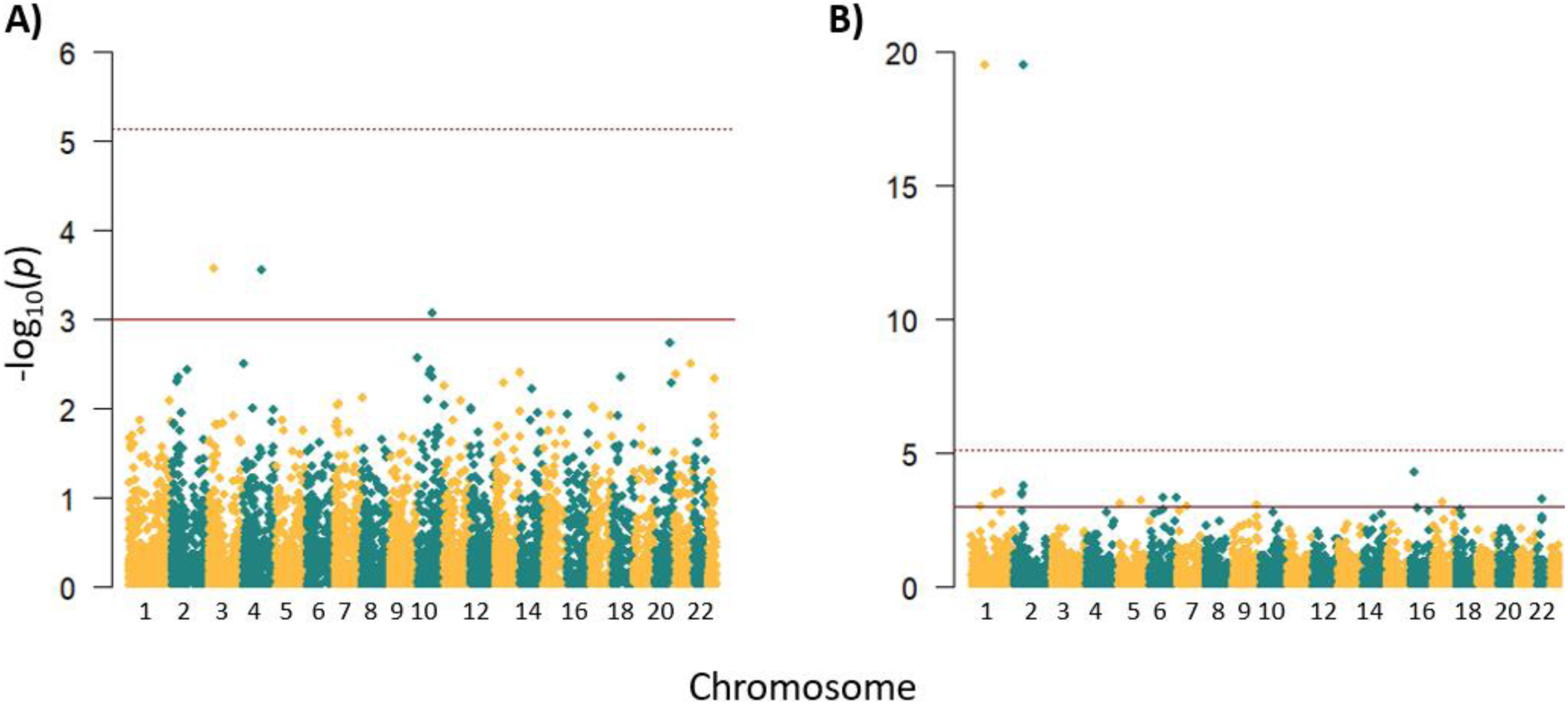
Manhattan plots of a MLM with DC as response including genetic structure as fixed effect and kinship and R as random effects **(A),** and a lfmm with annual precipitation as predictor of genotype values **(B)**. The solid and dashed lines represent a suggestive line defined at p < 0.001 and a Bonferroni corrected p < 0.05, respectively. Two association outliers for the GEA of annual precipitation are shown on chromosomes 1 and 2 **(B)**.

### Genotype-environment associations

We observed that most environmental variables considered were strongly correlated and deviated from normality (Fig. S8). Thus, we performed principal component analyses of these variables and used up to three PCs as synthetic environmental predictors in addition to testing the predictors mean annual precipitation, annual mean temperature, and altitude singularly. In lfmm, we discovered two strong outliers on chromosomes 1 and 2 associated with environmental variables modeled as synthetic PC axes (PC1, Fig. S9A, and PC2, Fig. S9B) that showed the strongest association with the annual precipitation (Fig.3B, S9C, Table S4) and the mean annual temperature range (Fig. S9D), but not with the altitude for which we found an associated outlier on chromosome 14 (Fig. S9E). Interestingly, the outliers associated with the PC axes of environmental predictors were also previously detected as associated with ADB damage by Stocks et al. (2019). The proteins affected by the outlier SNPs associated with the environmental variables are reported in Tables S4.

Two RDA models tested were overall significant (ANOVA p = 0.001), as were the constrained axes (ANOVA p = 0.001). We detected 21 candidate SNPs when using the first PC of environmental variables as synthetic predictor (Fig. 4), while 14 candidate SNPs were found associated with the predictor mean temperature annual range (Fig. S10A), 24 candidate SNPs with the predictor mean annual precipitation (Fig. S10A), and 20 candidate SNPs with the predictor altitude (Fig. S10B). These outliers are reported in Table S5 together with their position relative to genes and functional annotation. We observed that a majority of RDA outliers containing genes were reported to play a role in plant immune responses, including host-fungus interactions such as quinone-oxidoreductases (Heyno et al., 2013), abiotic stress responses, as well as epigenetic and gene expression regulation.

**Figure 4.**
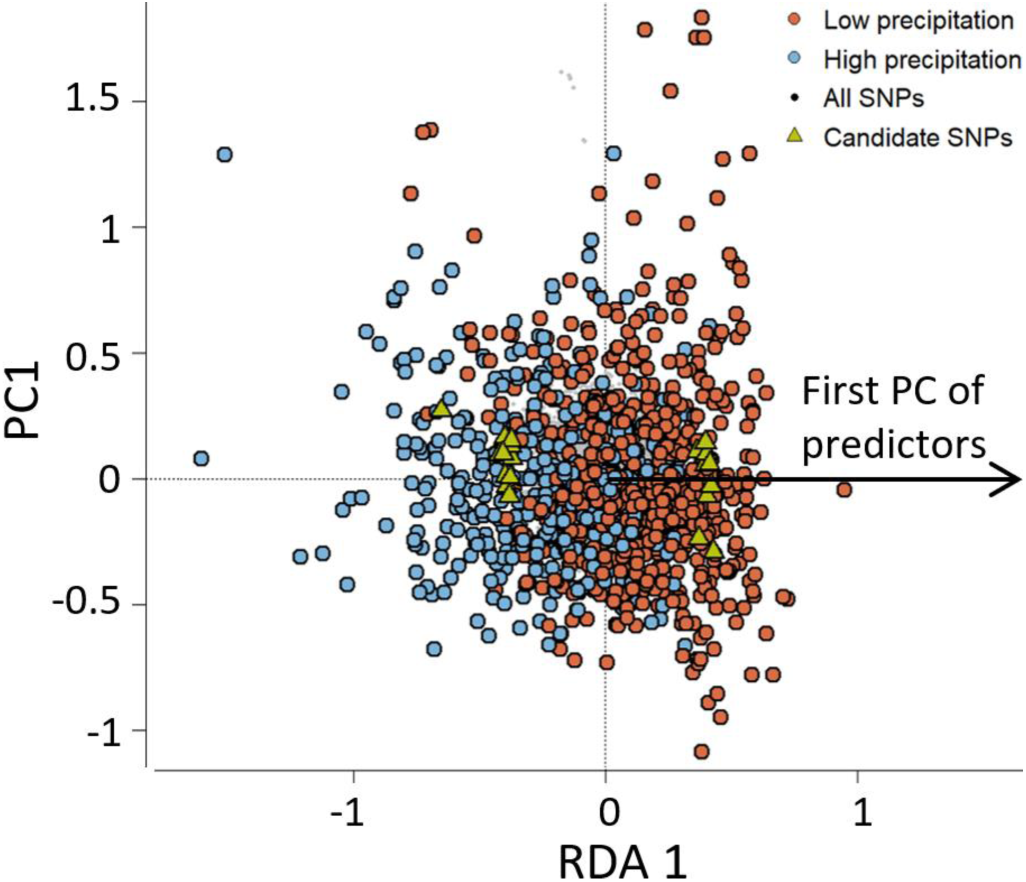
RDA plot showing the synthetic predictor of the environmental variables mean annual precipitation, mean annual temperature, and altitude. Candidate SNPs found along the RDA axis are depicted as green triangles, while the light blue and orange colors show a separation between individuals growing on sites with annual precipitation below (orange) or above (light blue) the average value.

## Discussion

Our endeavor to gather seeds from tolerant genotypes dispersed throughout Austria, followed by cultivation and a three-year monitoring period of tolerance to ADB, represents one of the largest efforts to date towards characterization of ADB tolerance of common ash at a national scale. This concerted effort also stands as one of the largest ash germplasm collections aimed at initiating a breeding program focused on enhancing tolerance to this detrimental disease. Such endeavors are underscored by scholarly consensus regarding the crucial role of breeding programs in addressing the challenges posed by ADB and bolstering forest resilience against its detrimental effects (Pautasso et al., 2013). Indeed, programmes to breed common ash trees with high ADB tolerance have been initiated in several European countries (McKinney et al. 2014; Enderle et al., 2017; Kjær et al., 2017; Pliūra et al., 2017; Marçais et al., 2022). In Austria, the conservation and breeding program “Ash in distress” was started in 2015 to provide reproductive material of common ash with high ADB tolerance for forestry and restoration purposes (Unger et al., 2021).

Previous research has undertaken GWAS of pooled whole genome data of common ash, with the overarching objective of pinpointing SNPs associated with ash dieback tolerance (Stocks et al., 2019; Doonan et al., 2023; Meger et al., 2024). The ultimate goal was to leverage the top candidates for subsequent screening of disease-tolerant common ash genotypes, as demonstrated by the development of the 4TREE Array (Guilbaoud et al., *in prep*). However, our results reveal an evident challenge in identifying a ready-to-use subset of genetic loci suitable for screening ash populations across Europe. This difficulty likely stems from the inherently polygenic nature of the trait under investigation that has been observed previously (McKinney et al., 2024; Stocks et al., 2019; Metheringham et al., 2022; Doonan et al., 2023). Moreover, the very rapid decay of LD observed in our data, also consistent with previous long-range LD estimates (Sollars et al., 2017), provides an explanation for the challenges encountered in capturing significant associated SNPs using the array. The fast LD decay implies that nearby genetic variants become less correlated over short distances, diminishing the power of GWAS to detect associations with SNPs in close proximity, leaving a majority of genomic regions uncovered (Fig. 2C). Our results thus underscore the importance of considering LD patterns during the design and establishment of genotyping tools, emphasizing the need for fine-mapping strategies to unravel complex genetic associations. An adaptive architecture characterized by many contributing loci of small effects (Barghi et al., 2020) and rapid LD decay are important limiting factors in association analyses using a small number of markers (Vos et al., 2017).

We observed that over 85 % of previously identified association outliers included in the 4TREE array (Stocks et al., 2019) were not informative in our sampling because they were monomorphic or characterized by rare alleles (i. e. MAF < 0.05). This observation highlights a significant disparity in genetic (potentially adaptive) diversity across different study areas. Such heterogeneity underscores the necessity for region-specific approaches in devising effective strategies aimed at bolstering tree tolerance to diseases. Consistently, other investigations on the tolerance to ADB found only low capacity of candidate markers from precedent studies to discriminate between putatively tolerant and susceptible genotypes (Menkis et al., 2019; Plumb, 2021), or found that SNPs sets showing high predictive accuracy on a training dataset failed to predict ADB-tolerance when tested on a different dataset (Meger et al., 2023).

Other studies have indeed highlighted the limitations of association studies in capturing the spectrum of adaptive genetic variation over large geographic scales (Long et al., 2013; Vilhjálmsson and Nordborg, 2013; Gloss et al., 2023). Their efficacy is limited by complicating factors such as genetic heterogeneity, allelic heterogeneity, rare variants, epistasis and population structure (Gloss et al., 2023), and likely also genetic redundancy in polygenic traits (Yeaman, 2022) driving differential defense responses across environments and geographic regions (Wohlmuth et al., 2018). The shallow population structure within our study area is unlikely to represent a major limiting factor, as expected for a tree species with long-range dispersal such as common ash (Heuertz et al., 2004; Sutherland et al., 2010; Beatty et al., 2015). Our results showed that rare alleles and allelic heterogeneity across the distribution range of common ash may more strongly impact the predictive power of SNPs arrays across national borders, suggesting that tolerance to ADB may be better captured through genetic marker panels developed covering larger geographic regions and multiple ecological conditions. Notably, our study area encompasses higher environmental heterogeneity than the geographic regions previously analyzed (i. e. by Stocks et al. 2019 in the case of the SNPs included in the array), potentially resulting in local adaptations involving different molecular pathways and genes.

Other significant challenges in the quest to identify causal SNPs likely lie in the sampling strategy and accuracy of the phenotyping in the present and previous studies. Our collection of susceptible and tolerant genotypes followed a first selection phase where mainly tolerant genotypes were selected (i. e. the mother trees selected in the field at the start of the study). This sampling strategy may have led to an underrepresentation of highly susceptible genotypes in our trial, possibly limiting the power to detect associations. Moreover, though ash dieback can lead to rapid mortality, it often progresses over many years; thus, the accuracy of categorizing trees as tolerant or susceptible increases with the number of years of observation. Although we expect most trees phenotyped as tolerant after three years to display durable tolerance, a portion of them may become diseased and succumb later, which would impact the results of our and past GWAS analyses.

An alternative approach to increase the accuracy of phenotyping is the artificial inoculation with *H. fraxineus* followed by the assessment of necrotic lesion length in the bark as an indicator of tolerance (but see also Mckinney et al., 2014, who raised concerns about the visible damage to the external woody parts as a reliable indicator of pathogen advancement and disease state). With artificial inoculations, the active defense of a genotype to limit the spread of *H. fraxineus* in plant tissues can be assessed; however, another known component of tolerance, i. e. the avoidance of the disease through early leaf senescence and yellowing, as well as probably early flushing, cannot be reliably estimated (McKinney et al. 2014). Molecular techniques, such as RNAseq and proteomics, and high-throughput phenotyping may also offer more nuanced insights into the complex interplay between genetic factors and tolerance to ADB. However, artificial inoculations and more sophisticated phenotyping will probably limit the number of seedlings/trees to be screened and might therefore be of limited use in large-scale selection programmes such as ours. A comprehensive understanding of infection and phenotyping limits is crucial for refining strategies in the pursuit of accurate identification of SNPs associated with tolerance to pathogens in tree populations.

Despite the inherent limitations in our dataset’s ability to capture association outliers linked to the degree of ADB damage, we successfully pinpointed loci associated with precipitation, altitude, and temperature gradients. These environmental factors appear to contribute significantly to the variation in damage caused by ADB (Chumanová et al., 2019; Grosdidier et al., 2020; Marçais et al., 2022), offering additional insights into the complex dynamics underlying disease intensity. Interestingly, the outliers detected in RDA analyses included a high number of genes putatively involved in immune and biotic stress in addition to abiotic stress responses, suggesting that the same genes involved in environmental responses bear the potential to influence tolerance to the pathogen. Our analyses also suggested that the flushing behavior influences the severity of ADB damage with late flushing individuals being more severely affected, which is in agreement with previous studies (McKinney et al. 2014).

The 4TREE array enabled us to effectively identify *F. excelsior* x *F. angustifolia* hybrid individuals and introgressants, a feature that holds promise for future applications. Leveraging this capability could lead to the development of SNP sets with high discriminatory power between *Fraxinus* species, enhancing our understanding of species interactions and aiding conservation efforts. Finally, we found two main genetic clusters within common ash corresponding to western and eastern Austria. Taken together, these results offer valuable knowledge for the creation of conservation and seed orchards of ADB-tolerant clones, which need to account for the environmental and geographic heterogeneity of common ash across Austria. For instance, two orchards were yet established at lower (eastern Upper Austria) and higher (eastern Tyrol) elevation sites using clones from sites below and above 600 m a.sl., respectively; moreover, one orchard was established in the province of Vorarlberg with clones from western Austria (Konrad et al., 2024).

In conclusion, our study sheds light on the challenges inherent in identifying a comprehensive set of genetic loci for screening ash populations across Europe, despite the advancements made in GWAS of ADB tolerance and the development of genotyping tools like the 4TREE array. We underscore the importance of considering LD patterns during the design of genotyping tools and highlight the need for fine-mapping strategies to unravel complex genetic associations effectively. Furthermore, we reveal the limited informativeness of previously identified association outliers in our sampling, advocating for region-specific approaches to enhance understanding of the genetic background of tree tolerance to diseases. Despite these challenges, our study demonstrates the utility of the 4TREE array in uncovering patterns of genetic structure and identifying *F. excelsior* x *F. angustifolia* hybrid individuals and its potential for using SNPs sets with high discriminatory power between ash species. Moreover, the array can be used to investigate how environmental factors such as precipitation and temperature gradients shape potentially adaptive genetic variation, providing valuable insights for future research and conservation efforts. Overall, these findings underscore the need for a comprehensive understanding of the challenges and opportunities in the pursuit of accurate identification of SNPs associated with pathogen tolerance in tree populations.

## Acknowledgements

This research was supported by “Ash in distress” (“Bedrohtes Erbgut Esche / Esche in Not”) projects (phase I: 2015-2019, phase II: 2019-2024) funded by the Austrian Chamber of Agriculture, the Federal Ministry of Agriculture, Forestry, Regions and Water Management (DaFNEplus grants nos. SNP and 101476), the State Forestry Administrations of all Federal Provinces and the Nature Conservation Departments of the Federal Provinces of Salzburg and Upper Austria, the Vienna City Administration (MA 49) and the Austrian Foresters Association. We thank Richard Buggs for insightful comments and feedback, Michael Stierschneider of AIT Tulln for DNA extractions, the team of gardeners of the BFW research nursery in Tulln for the germination and cultivation of the plants. Support in tree selection, seed harvest and ADB assessment was provided by Wernfried Zainer, Andreas Fera, Michael Kober-Eberhardt, Lambert Weißenbacher, Dominik Lorenschitz, Julia Rode, Rudolf Lebenits, Wilfried Nebenführ, Anton Aigner and many others. We also thank the editor and reviewers for helpful comments.

## Author contributions

TG, HK, GMU, SS and TK conceived and designed the study. HK, GMU and TK led and carried out the selection of tolerant mother trees, the germination and cultivation of seedlings, the establishment of the field trial and the data collection. AS conducted bioinformatic and statistical analyses of the data. AS plotted and interpreted the results with support by CT-M. AS wrote the manuscript and all authors revised and approved the manuscript.

## References

Andrić, I., Poljak, I., Milotić, M., Idžojtić, M., and Kajba, D. (2016). Leaf phenology characteristics of narrow-leaved ash (*Fraxinus angustifolia* Vahl) in clonal seed orchard. Šumarski list, 140: 117–126. doi: 10.31298/sl.140.3-4.2

Barghi, N., Hermisson, J., and Schlötterer, C. (2020). Polygenic adaptation: a unifying framework to understand positive selection. Nat. Rev. Genet. 21, 769–781. doi: 10.1038/s41576-020-0250-z

Beatty, G. E., Brown, J. A., Cassidy, E. M., Finlay, C. M. V., McKendrick, L., Montgomery, W. I., et al. (2015). Lack of genetic structure and evidence for long-distance dispersal in ash (*Fraxinus excelsior*) populations under threat from an emergent fungal pathogen: implications for restorative planting. Tree Genet. Genomes 11. doi: 10.1007/s11295-015-0879-5

Boshier, D., and Stewart, J. (2005). How local is local? Identifying the scale of adaptive variation in ash (*Fraxinus excelsior* L.): results from the nursery. Forestry 78, 135–143. doi: 10.1093/forestry/cpi013

Bourret, V., Dionne, M., and Bernatchez, L. (2014). Detecting genotypic changes associated with selective mortality at sea in Atlantic salmon: polygenic multilocus analysis surpasses genome scan. Mol. Ecol. 23, 4444–4457. doi: 10.1111/mec.12798

Bradbury, P. J., Zhang, Z., Kroon, D. E., Casstevens, T. M., Ramdoss, Y., and Buckler, E. S. (2007). TASSEL: software for association mapping of complex traits in diverse samples. Bioinformatics 23, 2633–2635. doi: 10.1093/bioinformatics/btm308

Capblancq, T., Luu, K., Blum, M. G. B., and Bazin, E. (n.d.). Evaluation of redundancy analysis to identify signatures of local adaptation. doi: 10.1101/258988

Caye, K., Deist, T. M., Martins, H., Michel, O., and François, O. (2016). TESS3: fast inference of spatial population structure and genome scans for selection. Mol. Ecol. Resour. 16, 540–548.

Caye, K., Jumentier, B., Lepeule, J., and François, O. (2019). LFMM 2: Fast and Accurate Inference of Gene-Environment Associations in Genome-Wide Studies. Mol. Biol. Evol. 36, 852–860.

Chaudhary, R., Rönneburg, T., Stein Åslund, M., Lundén, K., Durling, M. B., Ihrmark, K., et al. (2020). Marker-trait associations for tolerance to ash dieback in common ash (*Fraxinus excelsior* L.). For. Trees Livelihoods 11, 1083. doi: 10.3390/f11101083

Chumanová, E., Romportl, D., Havrdová, L., Zahradník, D., Pešková, V., and Černý, K. (2019). Predicting ash dieback severity and environmental suitability for the disease in forest stands. Scand. J. For. Res. 34, 254–266. doi: 10.1080/02827581.2019.1584638

Coker, T. L. R., Rozsypálek, J., Edwards, A., Harwood, T. P., Butfoy, L., and Buggs, R. J. A. (2019). Estimating mortality rates of European ash (*Fraxinus excelsior*) under the ash dieback (*Hymenoscyphus fraxineus*) epidemic. Plants, People, Planet 1, 48–58. doi: 10.1002/ppp3.11

Danecek, P., Auton, A., Abecasis, G., Albers, C. A., Banks, E., DePristo, M. A., et al. (2011). The variant call format and VCFtools. Bioinformatics 27, 2156–2158.

Danecek, P., Bonfield, J. K., Liddle, J., Marshall, J., Ohan, V., Pollard, M. O., et al. (2021). Twelve years of SAMtools and BCFtools. Gigascience 10, giab008.

Díaz-Yáñez, O., Mola-Yudego, B., Timmermann, V., Tollefsrud, M. M., Hietala, A. M., and Oliva, J. (2020). The invasive forest pathogen *Hymenoscyphus fraxineus* boosts mortality and triggers niche replacement of European ash (*Fraxinus excelsior)*. Sci. Rep. 10, 5310. doi: 10.1038/s41598-020-61990-4

Dobrowolska, D., Hein, S., Oosterbaan, A., Wagner, S., Clark, J., and Skovsgaard, J. P. (2011). A review of European ash (*Fraxinus excelsior* L.): implications for silviculture. Forestry 84, 133–148.

Dong, W., Li, E., Liu, Y., Xu, C., Wang, Y., Liu, K., et al. (2022). Phylogenomic approaches untangle early divergences and complex diversifications of the olive plant family. BMC Biol. 20, 92.

Doonan, J. M., Budde, K. B., Kosawang, C., Lobo, A., Verbylaite, R., Brealey, J. C., et al. (2023). Multiple, single trait GWAS and supervised machine learning reveal the genetic architecture of *Fraxinus excelsior* tolerance to ash dieback in Europe. bioRxiv. doi: 10.1101/2023.12.11.570802

Dray, S., Pélissier, R., Couteron, P., Fortin, M.-J., Legendre, P., Peres-Neto, P. R., et al. (2012). Community ecology in the age of multivariate multiscale spatial analysis. Ecological Monographs 82, 257–275. doi: 10.1890/11-1183.1

Dyderski, M. K., Paź, S., Frelich, L. E., and Jagodziński, A. M. (2018). How much does climate change threaten European forest tree species distributions? Glob. Chang. Biol. 24, 1150–1163.

Endelman, J. B., and Jannink, J.-L. (2012). Shrinkage estimation of the realized relationship matrix. G3 2, 1405–1413.

Enderle, R., Fussi, B., Lenz, H. D., Langer, G., Nagel, R., and Metzler, B. (2017). Ash dieback in Germany: research on disease development, resistance and management options. In: Vasaitis, R., and Enderle, R. (editors), Dieback of European Ash (Fraxinus spp.) – Consequences and Guidelines for Sustainable Management. Report on European Cooperation in Science & Technology COST Action FP1103 FRAXBACK, SLU Service/Repro, Uppsala, 89–114.

Fernandez-Manjarres, J. F., Gerard, P. R., Dufour, J., Raquin, C., and Frascaria-Lacoste, N. (2006). Differential patterns of morphological and molecular hybridization between *Fraxinus excelsior* L. and *Fraxinus angustifolia* Vahl (Oleaceae) in eastern and western France. Mol. Ecol. 15, 3245–3257. doi: 10.1111/j.1365-294X.2006.02975.x

Forester, B. R., Lasky, J. R., Wagner, H. H., and Urban, D. L. (2018). Comparing methods for detecting multilocus adaptation with multivariate genotype-environment associations. Mol. Ecol. 27, 2215–2233.

Freinschlag, C. (2013). Untersuchungen zum Eschentriebsterben in Eschen-Samenplantagen in Österreich [Investigations on ash dieback in ash seed plantations in Austria]. Master thesis, Institute of Forest Entomology, Forest Pathology and Forest Protection (IFFF), University of Natural Resources and Life Sciences, Vienna (BOKU), Vienna, Austria, 140 pp (in German).

Frichot, E., Schoville, S. D., Bouchard, G., and François, O. (2013). Testing for associations between loci and environmental gradients using latent factor mixed models. Mol. Biol. Evol. 30, 1687C1699.

Friedman, J., Hastie, T., and Tibshirani, R. (2010). Regularization paths for generalized linear models via coordinate descent. J. Stat.Softw. 33, 1–22.

Friendly, M., and Meyer, D. (2015). Discrete Data Analysis with R: Visualization and Modeling Techniques for Categorical and Count Data. CRC Press.

George, J.-P., Sanders, T. G. M., Timmermann, V., Potočić, N., and Lang, M. (2022). European-wide forest monitoring substantiates the necessity for a joint conservation strategy to rescue European ash species (*Fraxinus* spp.). Sci. Rep. 12, 4764. doi: 10.1038/s41598-022-08825-6

Gloss, A. D., Steiner, M. C., Novembre, J., and Bergelson, J. (2023). The design of mapping populations: Impacts of geographic scale on genetic architecture and mapping efficacy for defense and immunity. Curr. Opin. Plant Biol. 74, 102399.

Goberville, E., Hautekèete, N.-C., Kirby, R. R., Piquot, Y., Luczak, C., and Beaugrand, G. (2016). Climate change and the ash dieback crisis. Sci. Rep. 6, 35303. doi: 10.1038/srep35303

Grosdidier, M., Scordia, T., Ioos, R., and Marçais, B. (2020). Landscape epidemiology of ash dieback. J. Ecol. 108, 1789–1799.

Gross, A., Holdenrieder, O., Pautasso, M., Queloz, V., Sieber, T.N. (2014). *Hymenoscyphus pseudoalbidus*, the causal agent of European ash dieback. Mol. Plant Pathol. 15, 5–21.

Guilbaud, R., Biselli, C., Buiteveld, J., Cattivelli, L., Copini, P., Dowkiw, A., et al. (in prep). Development of a new tool (4TREE) for adapted genome selection in European tree species..

Guilbaud, R., Biselli, C., Buiteveld, J., Cattivelli, L., Copini, P., Dowkiw, A., et al. (2020). Development of a new tool (4TREE) for adapted genome selection in European tree species. Proceedings of the Gentree Symposium, (Avignon).

Harper, A. L., McKinney, L. V., Nielsen, L. R., Havlickova, L., Li, Y., Trick, M., et al. (2016). Molecular markers for tolerance of European ash (*Fraxinus excelsior*) to dieback disease identified using Associative Transcriptomics. Sci. Rep. 6, 19335.

Heinze, B., Tiefenbacher, R., Litschauer, R., and Kirisits, T. (2017): Ash dieback in Austria – history, current situation and outlook. In: Vasaitis, R., and Enderle, R. (editors), Dieback of European Ash (Fraxinus spp.) – Consequences and Guidelines for Sustainable Management. SLU Service/Repro, Uppsala, ISBN 978-91-576-8696-1, p. 33–52.

Heuertz, M., Hausman, J.-F., Hardy, O. J., Vendramin, G. G., Frascaria-Lacoste, N., and Vekemans, X. (2004). Nuclear microsatellites reveal contrasting patterns of genetic structure between western and southeastern European populations of the common ash (*Fraxinus excelsior* L.). Evolution 58, 976–988.

Heyno, E., Alkan, N., and Fluhr, R. (2013). A dual role for plant quinone reductases in host-fungus interaction. Physiol. Plant. 149, 340–353.

Huff, M., Seaman, J., Wu, D., Zhebentyayeva, T., Kelly, L. J., Faridi, N., et al. (2022). A high-quality reference genome for *Fraxinus pennsylvanica* for ash species restoration and research. Mol. Ecol. Resour. 22, 1284–1302.

Kirisits, T. and Freinschlag, C. (2012). Ash dieback caused by *Hymenoscyphus pseudoalbidus* in a seed plantation of *Fraxinus excelsior* in Austria. J. Agric. Ext. Rural Dev. 4, 184–191. doi: 10.5897/jaerd12.046

Kjær, E. D., McKinney, L. V., Nielsen, L. R., Hansen, L. N., and Hansen, J. K. (2012). Adaptive potential of ash (*Fraxinus excelsior*) populations against the novel emerging pathogen *Hymenoscyphus pseudoalbidus*. Evol. Appl. 5, 219–228. doi: 10.1111/j.1752-4571.2011.00222.x

Kjær, E. D., McKinney, L. V., Hansen, L. N., Olrik, D. C., Lobo, A., Thomsen, I., Hansen, J. K., and Nielsen, L. R. (2017). Genetics of ash dieback resistance in a restoration context – experiences from Denmark. In: Vasaitis, R., and Enderle, R. (editors), Dieback of European ash (Fraxinus spp.) – Consequences and Guidelines for Sustainable Management. Report on European Cooperation in Science & Technology COST Action FP1103 FRAXBACK, SLU Service/Repro, Uppsala, 106–114.

Konrad, H., Szukala, A., Aigner, A., Schueler, S., Schwanda, K., Neidel, V., Kirisits, T., and Unger, G. M. (2024). Die Esche kehrt zurück. Forstzeitung, 07-2024, 26–28.

Landolt, J., Gross, A., Holdenrieder, O., and Pautasso, M. (2016). Ash dieback due to *Hymenoscyphus fraxineus*: what can be learnt from evolutionary ecology? Plant Pathol. 65, 1056–1070. doi: 10.1111/ppa.12539

Langmead, B., and Salzberg, S. L. (2012). Fast gapped-read alignment with Bowtie 2. Nat. Methods 9, 357–359. doi: 10.1038/nmeth.1923

Legendre, P., Oksanen, J., and ter Braak, C. J. F. (2011). Testing the significance of canonical axes in redundancy analysis. Methods in Ecology and Evolution 2, 269–277. doi: 10.1111/j.2041-210x.2010.00078.x

Lobo, A., McKinney, L. V., Hansen, J. K., Kjaer, E. D., and Nielsen, L. R. (2015). Genetic variation in dieback resistance in *Fraxinus excelsior* confirmed by progeny inoculation assay. For. Pathol. 45, 379–387. doi: 10.1111/efp.12179

Long, Q., Rabanal, F. A., Meng, D., Huber, C. D., Farlow, A., Platzer, A., et al. (2013). Massive genomic variation and strong selection in *Arabidopsis thaliana* lines from Sweden. Nat. Genet. 45, 884–890.

Madsen, C. L., Kosawang, C., Thomsen, I. M., Hansen, L. N., Nielsen, L. R., and Kjaer, E. D. (2021). Combined progress in symptoms caused by *Hymenoscyphus fraxineus* and *Armillaria* species, and corresponding mortality in young and old ash trees. For. Ecol. Manag. 491, 119177. doi: 10.1016/j.foreco.2021.119177

Marçais, B., Husson, C., Caël, O., Dowkiw, A., Saintonge, F.-X., Delahaye, L., et al. (2017). Estimation of ash mortality induced by *Hymenoscyphus fraxineus* in France and Belgium. Balt. For. 23, 159–167.

Marçais, B., Kosawang, C., Laubray, S., Kjær, E., and Kirisits, T. (2022). Chapter 13 - Ash dieback, In: Kovalchuk, A. F., (editor), Forest Microbiology Volume 2: Forest Tree Health. Academic Press, 2022. p. 215–237. ISBN 0323984487, 9780323984485. doi: 10.1016/B978-0-323-85042-1.00022-7

Marçais, B., Giraudel, A., and Husson, C. (2023). Ability of the ash dieback pathogen to reproduce and to induce damage on its host are controlled by different environmental parameters. PLoS Pathog. 19, e1010558. doi: 10.3390/f12030340

Matisone, I., Matisons, R., and Jansons, Ā. (2021). The struggle of ash—insights from long-term survey in Latvia. Forests 12, 340.

McKinney, L. V., Nielsen, L. R., Collinge, D. B., Thomsen, I. M., Hansen, J. K., and Kjaer, E. D. (2014). The ash dieback crisis: genetic variation in resistance can prove a long-term solution. Plant Pathol. 63, 485–499. doi: 10.1111/ppa.12196

McMullan, M., Rafiqi, M., Kaithakottil, G., Clavijo, B. J., Bilham, L., Orton, E., et al. (2018). The ash dieback invasion of Europe was founded by two genetically divergent individuals. Nat Ecol Evol 2, 1000–1008. doi: 10.1038/s41559-018-0548-9

Meger, J., Ulaszewski, B., Pałucka, M, Kozioł, C., and Burczyk, J. (2024). Genomic prediction of resistance to *Hymenoscyphus fraxineus* in common ash (*Fraxinus excelsior* L.) populations. Evol. Appl. 17: e13694, doi: 10.1111/eva.13694

Menkis, A., Bakys, R., Åslund, M.S., Davydenko, K., Elfstrand, M., Stenlid, J., Vasaitis, R. (2019). Identifying *Fraxinus excelsior* tolerant to ash dieback: Visual field monitoring versus a molecular marker approach. For. Pathol. 2020;50:e12572. doi: 10.1111/efp.12572

Metheringham, C. L., Plumb, W. J., Stocks, J. J., Kelly, L. J., Gorriz, M. N., Moat, J., et al. (2022). Rapid polygenic adaptation in a wild population of ash trees under a novel fungal epidemic. bioRxiv. doi: 10.1101/2022.08.01.502033

Mitchell, R. J., Beaton, J. K., Bellamy, P. E., Broome, A., Chetcuti, J., Eaton, S., et al. (2014). Ash dieback in the UK: A review of the ecological and conservation implications and potential management options. Biol. Conserv. 175, 95–109. doi: 10.1016/j.biocon.2014.04.019

Oksanen, J., Blanchet, F. G., Kindt, R., Legendre, P., Minchin, P. R., O’Hara, R. B., et al. (2012). vegan: Community Ecology Package. Software http://CRAN.R-project.org/package=vegan

Pautasso, M., Aas, G., Queloz, V., and Holdenrieder, O. (2013). European ash (*Fraxinus excelsior*) dieback – A conservation biology challenge. Biol. Conserv. 158, 37–49. doi: 10.1016/j.biocon.2012.08.026

Pliūra, A., Lygis, V., Marčiulyniene, D., Suchockas, V., and Bakys, R. (2015). Genetic variation of *Fraxinus excelsior* half-sib families in response to ash dieback disease following simulated spring frost and summer drought treatments. iForest 9: 12–22. doi: 10.3832/ifor1514-008

Pliūra, A., Bakys, R., Suchockas, V., Marčiulynienė, G., Verbyla, V., and Lygis, V. (2017). Ash dieback in Lithuania: disease history, research on impact and genetic variation in disease resistance, tree breeding and options for forest management. In: Vasaitis, R., and Enderle, R. (editors), Dieback of European Ash (Fraxinus spp.) – Consequences and Guidelines for Sustainable Management. Report on European Cooperation in Science & Technology COST Action FP1103 FRAXBACK, SLU Service/Repro, Uppsala, 89–114.

Plumb, W. J. (2021). Resources for the development of ash trees resistant to ash dieback. PhD diss., Queen Mary University of London. uri: https://qmro.qmul.ac.uk/xmlui/handle/123456789/83361

Rellstab, C., Gugerli, F., Eckert, A. J., Hancock, A. M., and Holderegger, R. (2015). A practical guide to environmental association analysis in landscape genomics. Mol. Ecol. 24, 4348–4370. doi: 10.1111/mec.13322

Verdugo, R. A., and Orostica, K. Y. (2017). chromPlot: Global visualization tool of genomic data. Bioconductor. doi: 10.18129/B9.bioc.chromPlot

Vilhjálmsson, B., and Nordborg, M. (2013) The nature of confounding in genome-wide association studies. Nat Rev Genet 14, 1–2. doi: 10.1038/nrg3382

Saumitou-Laprade, P., Vernet, P., Dowkiw, A., Bertrand, S., Billiard, S., Albert, B., et al. (2018). Polygamy or subdioecy? The impact of diallelic self-incompatibility on the sexual system in *Fraxinus excelsior* (Oleaceae). Proc. Biol. Sci. 285. doi: 10.1098/rspb.2018.0004

Sollars, E. S. A., and Buggs, R. J. A. (2018). Genome-wide epigenetic variation among ash trees differing in susceptibility to a fungal disease. BMC Genomics 19, 502. doi: 10.1186/s12864-018-4874-8

Sollars, E. S. A., Harper, A. L., Kelly, L. J., Sambles, C. M., Ramirez-Gonzalez, R. H., Swarbreck, D., et al. (2017). Genome sequence and genetic diversity of European ash trees. Nature 541, 212–216. doi: 10.1038/nature20786

Stocks, J. J., Metheringham, C. L., Plumb, W., Lee, S. J., Kelly, L. J., Nichols, R. A., and Buggs, R. J. A. (2019). Genomic basis of European ash tree resistance to ash dieback fungus. *Nat*. Ecol. Evol. 3: 1686–1696. doi: 10.1101/626234

Sutherland, B. G., Belaj, A., Nier, S., Cottrell, J. E., Vaughan, S. P., Hubert, J., and Russell, K. (2010). Molecular biodiversity and population structure in common ash (*Fraxinus excelsior* L.) in Britain: implications for conservation. Mol. Ecol. 19, 2196–2211. doi: 10.1111/j.1365-294X.2009.04376.x

Thomasset, M., Fernandez-Manjarrés, J. F., Douglas, G. C., Frascaria-Lacoste, N., Raquin, C., and Hodkinson, T. R. (2011). Molecular and morphological characterization of reciprocal F1 Hybrid ash (*Fraxinus excelsior*×*Fraxinus angustifolia*, Oleaceae) and parental species reveals asymmetric character inheritance. Int. J. Plant Sci. 172, 423–433. doi: 10.1086/658169

Turner, S. D.(2018). qqman: an R package for visualizing GWAS results using Q-Q and manhattan plots. J. of Open Source Softw., 3, 731, doi: 10.21105/joss.00731

Unger, G. M., Konrad, H., Schwanda, K., Cech, T. L., Hoch, G., Fera, A., Kirisits, T., and Geburek, T. (2021). Ash in distress: conservation and resistance breeding programme for *Fraxinus excelsior* in Austria. In: Sallmannshofer, M., Schüler, S., Westergren, M. (editors), Perspectives for forest and conservation management in riparian forests. Slovenian Forestry Institute, Silva Slovenica Publishing Centre, Ljubljana, 146–150. doi: 10.20315/SFS.169

Vasaitis, R., and Enderle, R. (2017). Dieback of European Ash (Fraxinus spp.): Consequences and Guidelines for Sustainable Management. Report on European Cooperation in Science & Technology COST Action FP1103 FRAXBACK, SLU Service/Repro, Uppsala, Sweden.

Verdugo, R. A., and Orostica, K. Y. (2024). chromPlot: Global visualization tool of genomic data. https://git.bioconductor.org/packages/chromPlot

Vos, P. G., Paulo, M. J., Voorrips, R. E., Visser, R. G. F., van Eck, H. J., and van Eeuwijk, F. A. (2017). Evaluation of LD decay and various LD-decay estimators in simulated and SNP-array data of tetraploid potato. Theor. Appl. Genet. 130, 123–135. doi: 10.1007/s00122-016-2798-8

Wohlmuth, A., Essl, F., and Heinze, B. (2018). Genetic analysis of inherited reduced susceptibility of *Fraxinus excelsior* L. seedlings in Austria to ash dieback. Forestry 91, 514–525. doi: 10.1093/forestry/cpy012

Yeaman, S. (2022). Evolution of polygenic traits under global vs local adaptation. Genetics 220. doi: 10.1093/genetics/iyab134

Zukrigl, K. (1997). Seltene Eschen I [Rare ash species I]. In: WWF Austria (editor), Proceedings of the symposium “Zukunft für gefährdete Baumarten? Rückbringung und Förderung seltener und gefährdeter Baum- und Straucharten” [“A future for endangered tree species? Restoration and promotion of rare and endangered tree and shrub species”], Vienna, Austria, 1 October 1997: 7-10 (in German).

