## supplementary_figures for "Limitations of *ad hoc* genotyping in detecting ash dieback tolerance in *Fraxinus excelsior*"

**A)**

| Damage Class [DC] | Class Range | Class Mean |
|---|---|---|
| 1 | 0 % (not diseased) | 0 % |
| 2 | up to 10 % | 5 % |
| 3 | > 10 % to 50 % | 30 % |
| 4 | > 50 % to 90 % | 70 % |
| 5 | > 90 % to < 100 % | 95 % |
| 6 | 100 % (damaged and dead) | 100 % |

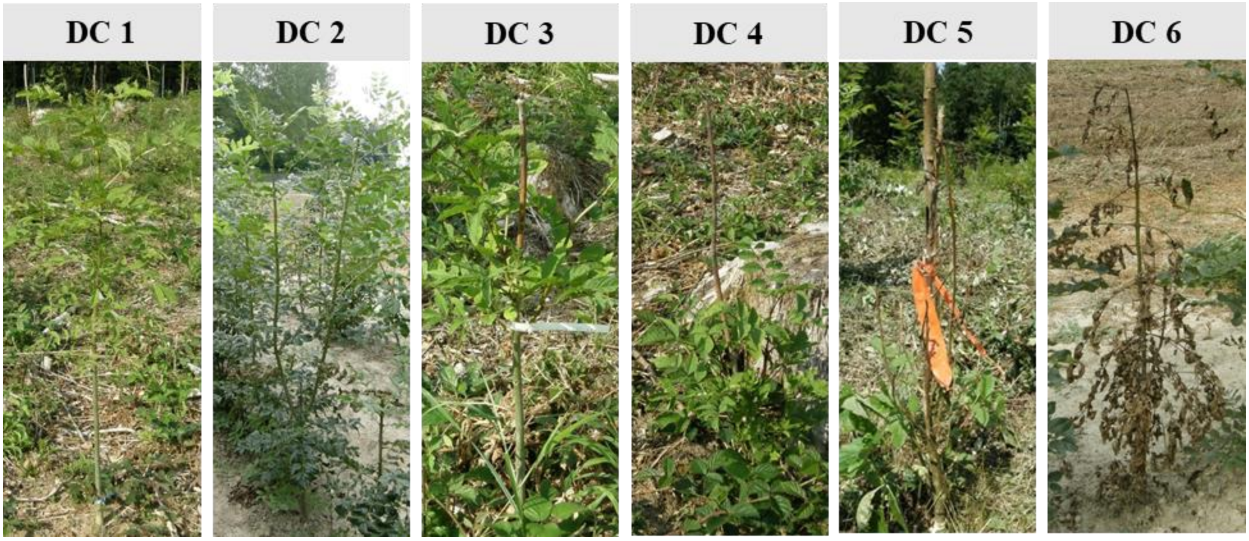


**B)**

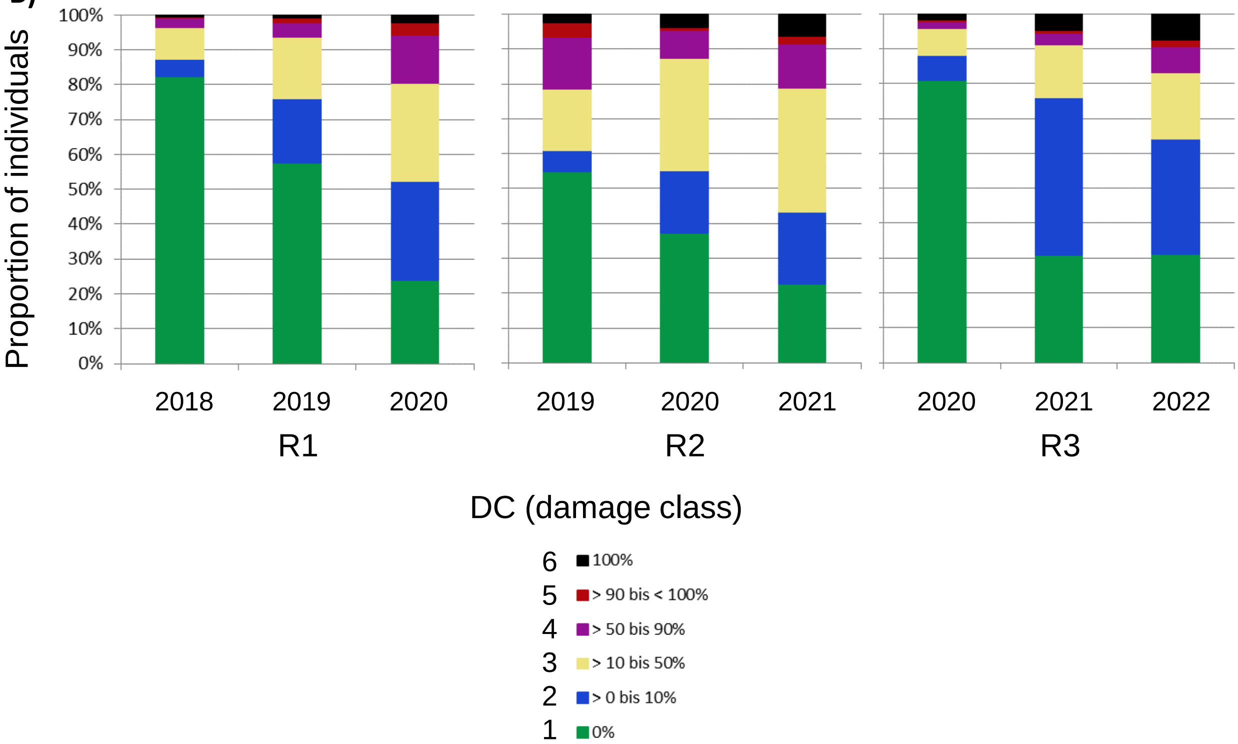

Fig. S1 Scheme to assess the damage on woody parts due to ash dieback used in the Austrian program “Ash in Distress” to breed ADB tolerant Fraxinus excelsior **(A)**. Proportion of individuals assigned to each one of six damage classes (DC) over three years of damage assessment in three field test sites of seedlings germinated in consecutive years (R1, R2, and R3) **(B)**. Photos in **(B)** are by Thomas Kirisits and were also used in Wohlmuth et al., 2018.

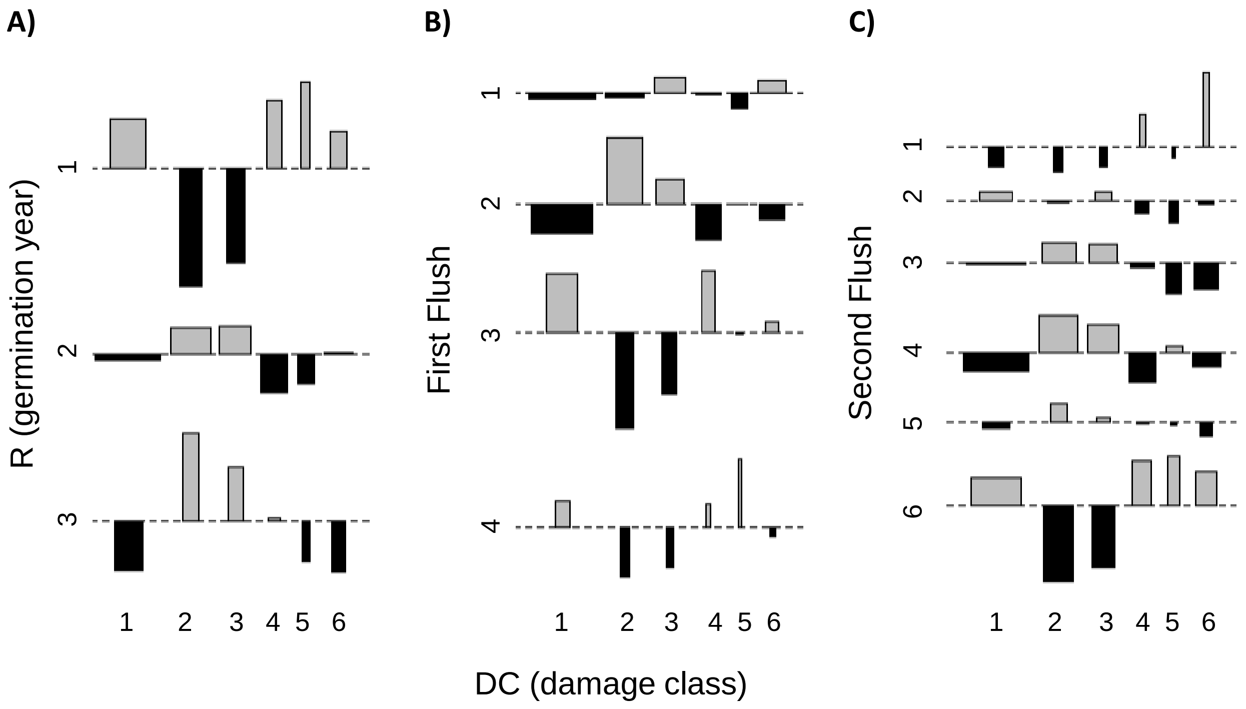


Fig. S2 Cohen-friendly association plots displaying a deviation from indipendence of DC (x-axis) and the variables R **(A)** and first **(B)** and second **(C)** Flush. The rectangles in each row are positioned relative to a baseline that indicates independence. If the frequency of individuals found in the cell of the underlying contigency table is greater or lower than the expectation under the hypothesis of independence, the rectangle rises above or falls below the baseline, respectively.

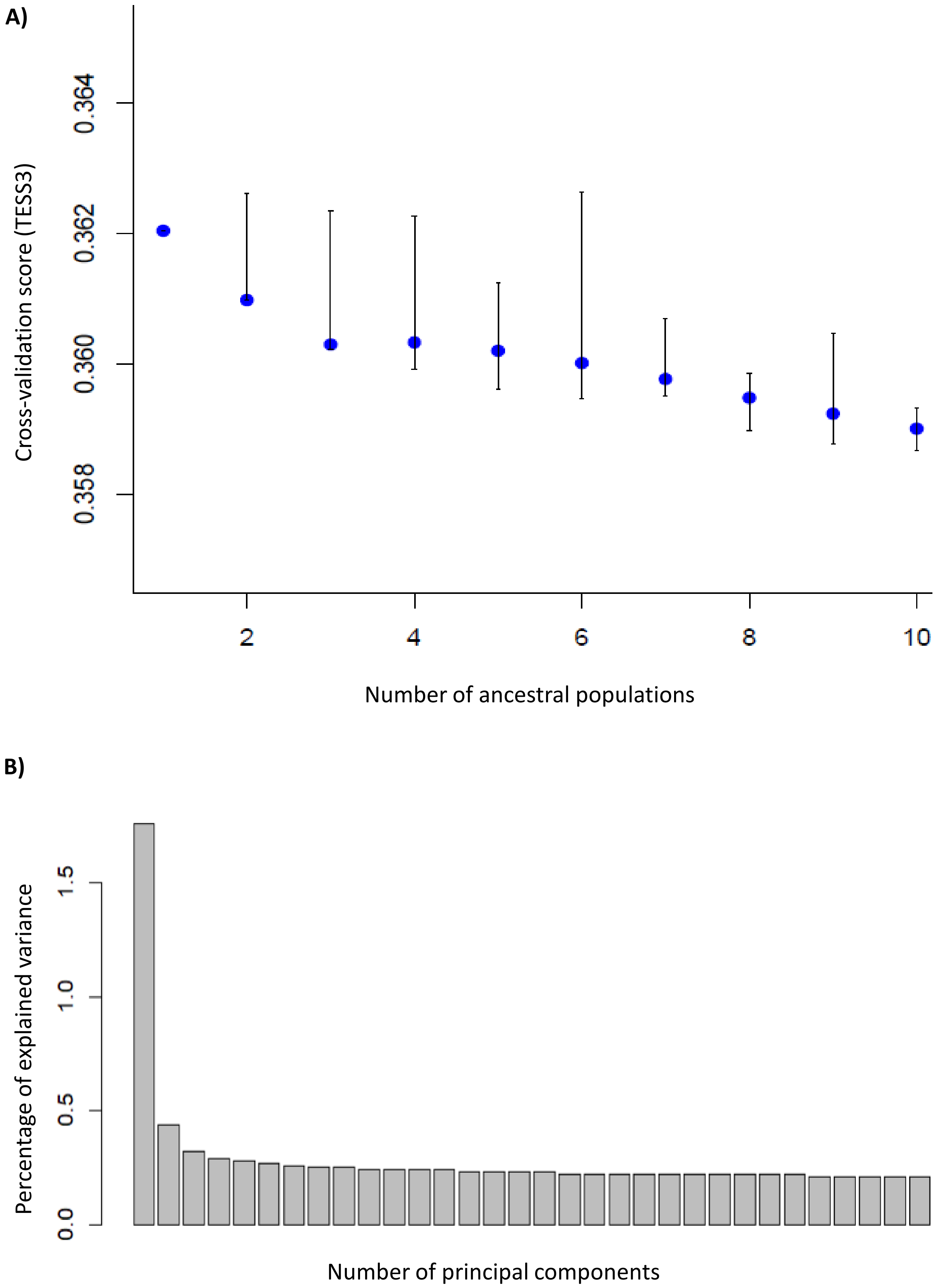
A)
Cross-validation score (TESS3)
0.358
0.360
0.362
0.364
2
4
6
8
10
Number of ancestral populations
B)
Percentage of explained variance
0.0
0.5
1.0
1.5
Number of principal components

Fig. S3 Cross-validation plot obtained in TESS3 for K from 1 to 10. The curve plateaus at K = 3, but finer levels of population structure can be detected from k >= 6 **(A)**. Percentage of variance explained by components 1 to 20 in PCA. Similarly to the TESS3 cross-validation scores, the increment of variance explained by components larger than 3 is very small **(B)**.

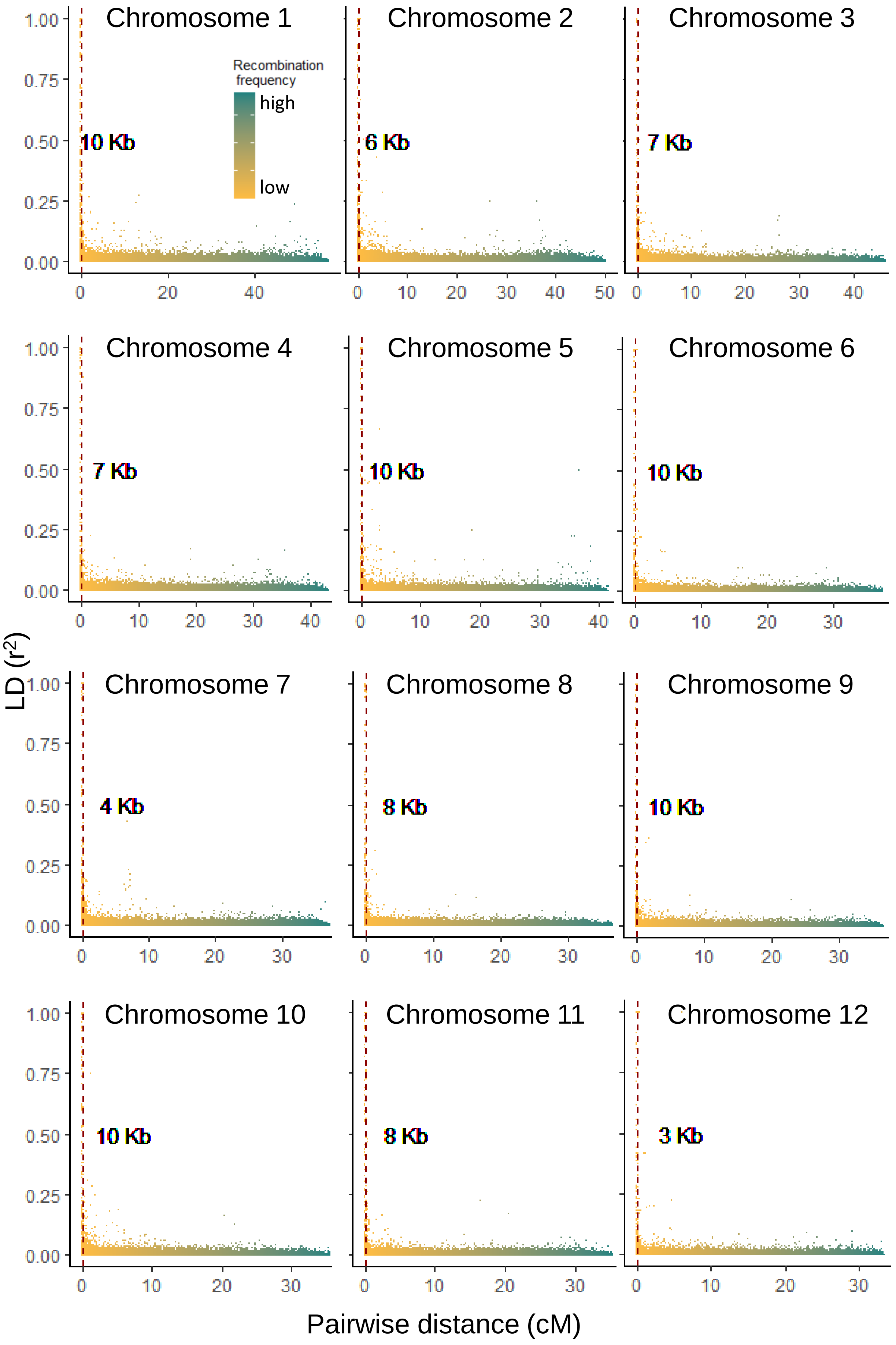

Chromosome 1
Chromosome 2
Chromosome 3
Chromosome 4
Chromosome 5
Chromosome 6
Chromosome 7
Chromosome 8
Chromosome 9
Chromosome 10
Chromosome 11
Chromosome 12
Recombination frequency
high
low
10 Kb
6 Kb
7 Kb
7 Kb
10 Kb
10 Kb
4 Kb
8 Kb
10 Kb
10 Kb
8 Kb
3 Kb
LD ($r^2$)
Pairwise distance (cM)

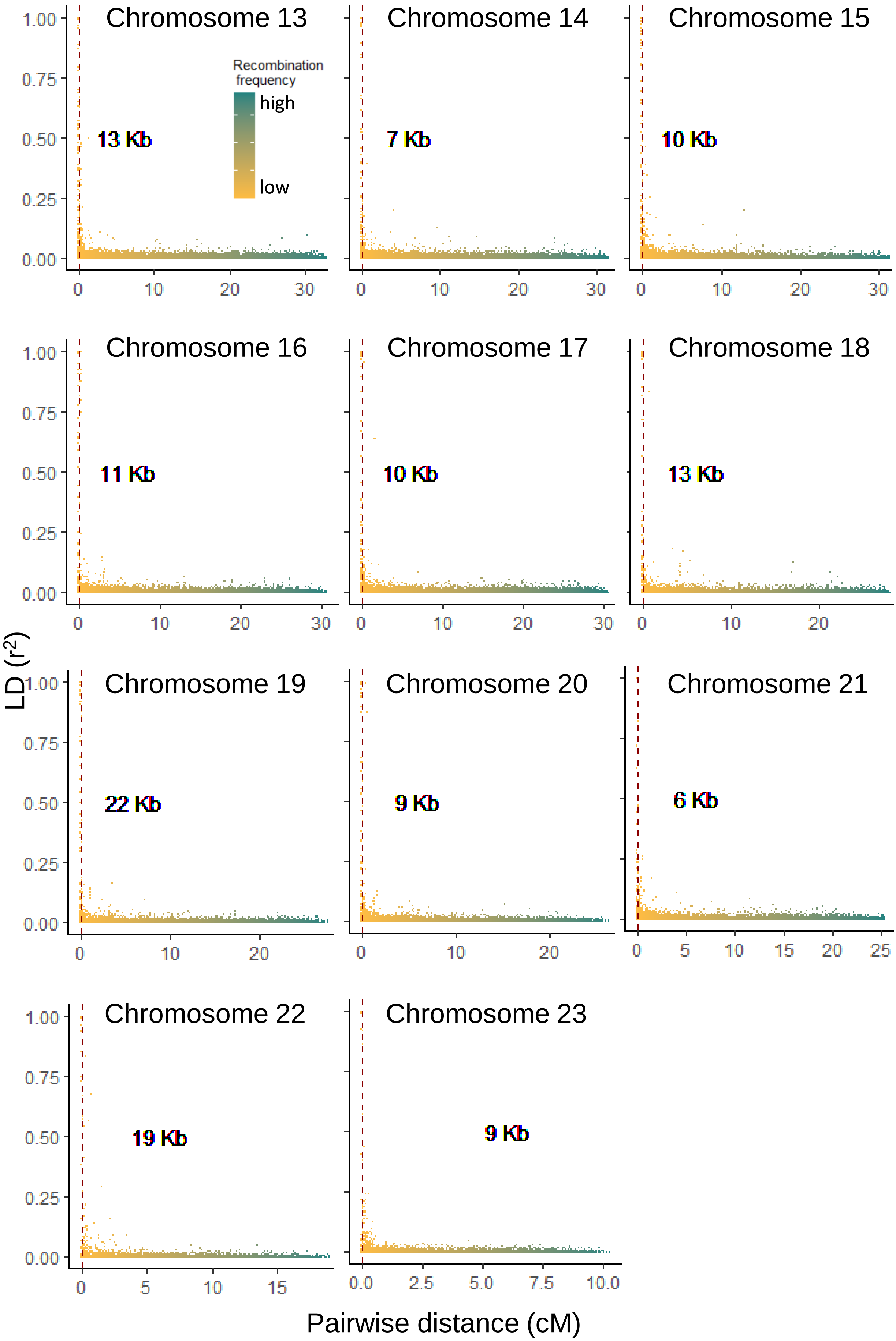

Chromosome 13
Recombination frequency
high
low
13 Kb
Chromosome 14
7 Kb
Chromosome 15
10 Kb
Chromosome 16
11 Kb
Chromosome 17
10 Kb
Chromosome 18
13 Kb
Chromosome 19
22 Kb
Chromosome 20
9 Kb
Chromosome 21
6 Kb
Chromosome 22
19 Kb
Chromosome 23
9 Kb
LD (r2)
Pairwise distance (cM)

Fig. S4 Rate of LD decay by chromosome. The color scale shows the recombination frequency for a specific genomic distance expressed in centiMorgan (cM). The vertical dashed line and value reported in bold indicate the distance at which LD falls to half of its maximum value.

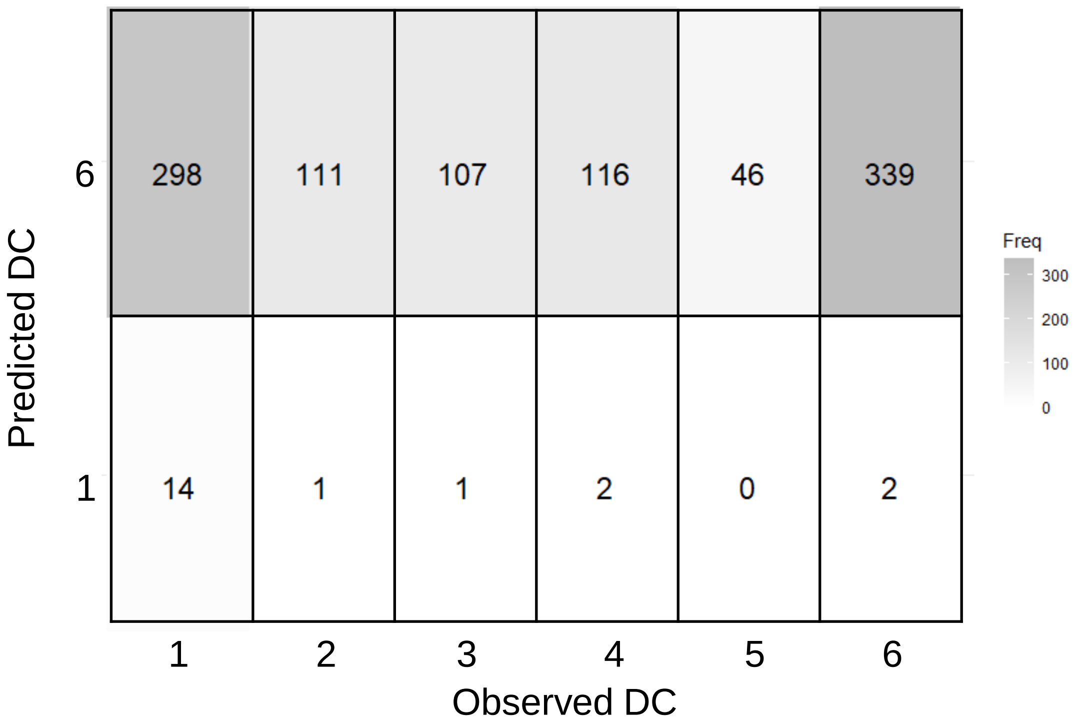


Fig. S5 Confusion matrix showing the match between predicted observed damage classes based on 13 SNPs found associated with the DC using multinomial logistic regression in glmnet.

A)

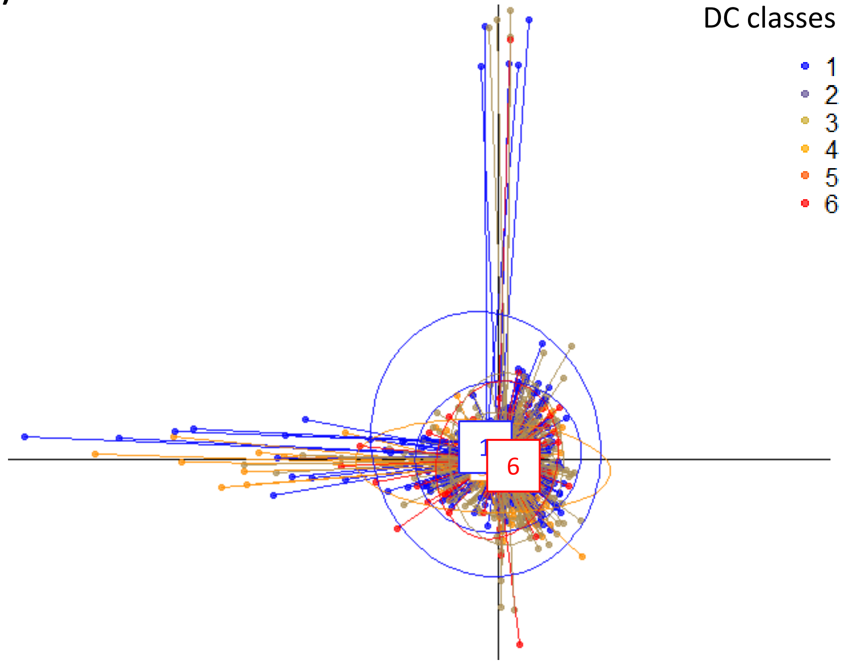


B)

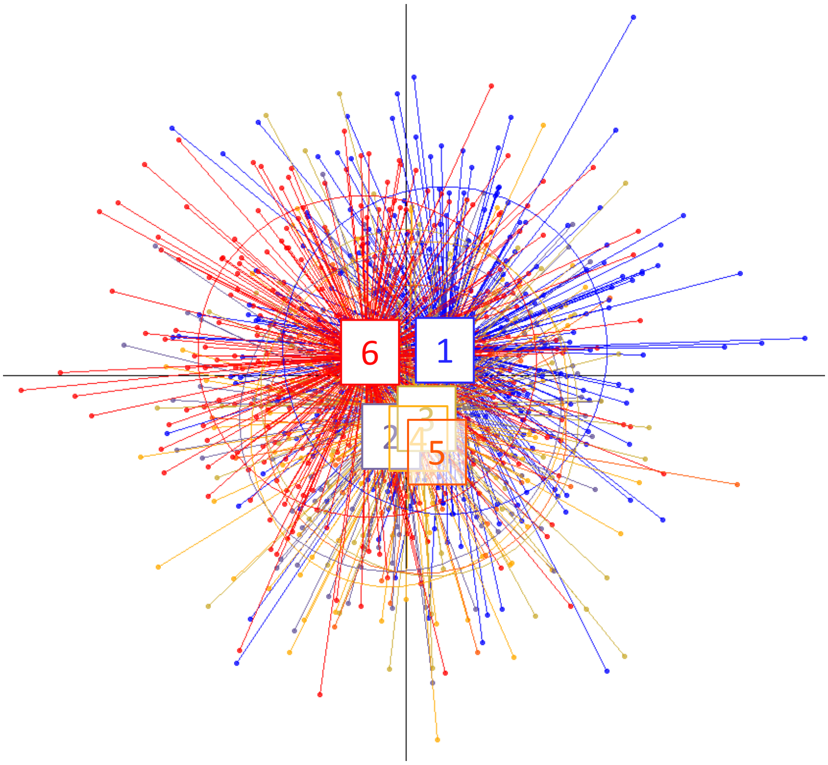


Fig. S6 DAPC cross-validation aimed at discriminating between the DC classes after accounting for genetic structure (i.e. using corrected genotypes). DAPC of all SNPs tested in the GWAS analyses showing a complete overlap between DC classes **(A)**. DAPC based on the 13 SNPs identified as associated with DC by multinomial logistic regression in glmnet failing to accurately discriminate between DC categories **(B)**.

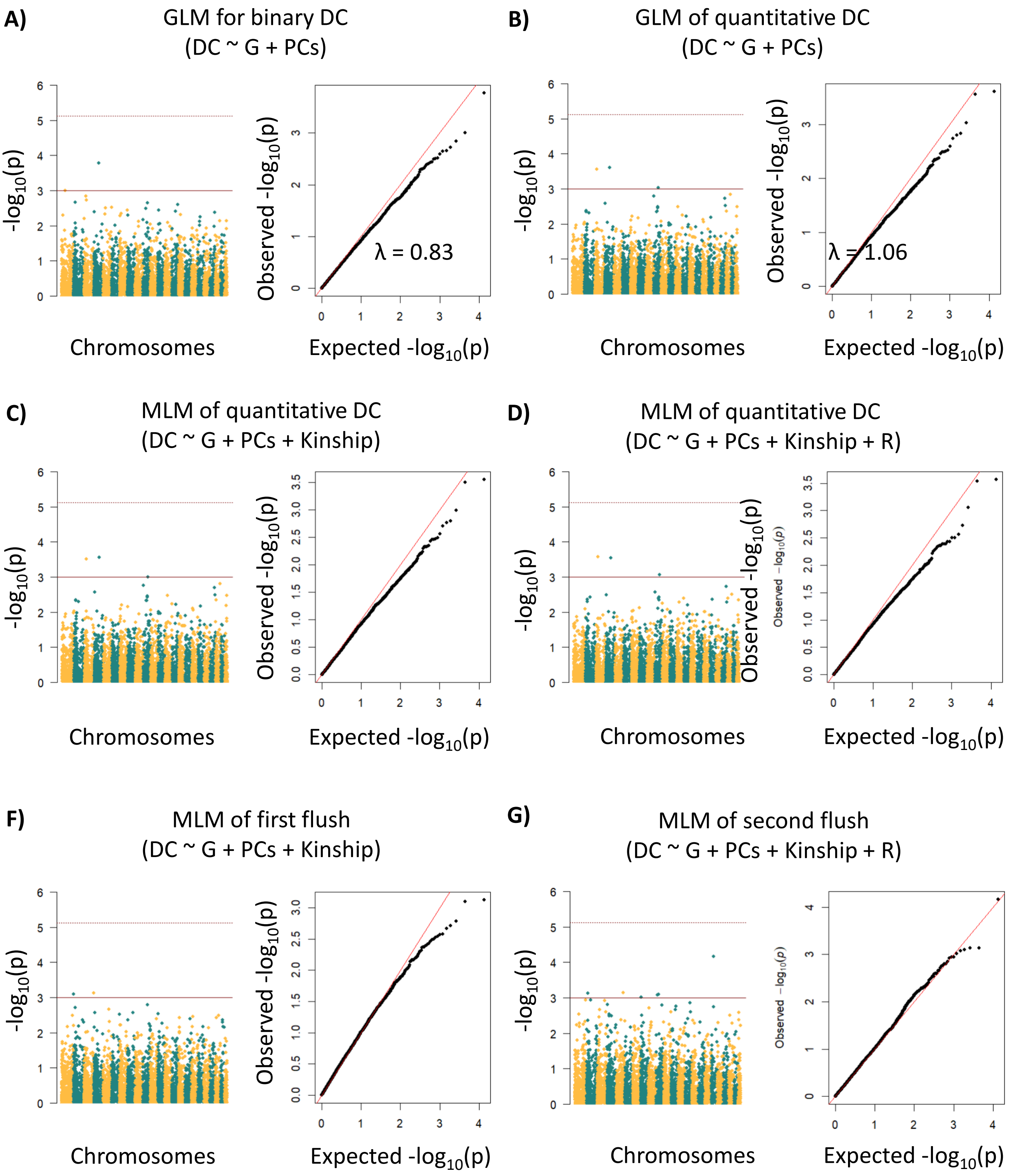


Fig. S7 Manhattan and QQ plots of GLM MLM with DC **(A-D)**, and first **(F)** and second **(G)** flushing as response variables. Models used are reported in the title of each pair of plots. The solid and dashed lines in Manhattan plots represent a suggestive line defined at $p < 0.001$ and a Bonferroni corrected $p < 0.05$, respectively. G = genotype matrix; DC = damage classe; PCs = principal components used as proxies for genetic structure; R = R test;2nd Flush = Second flushing.

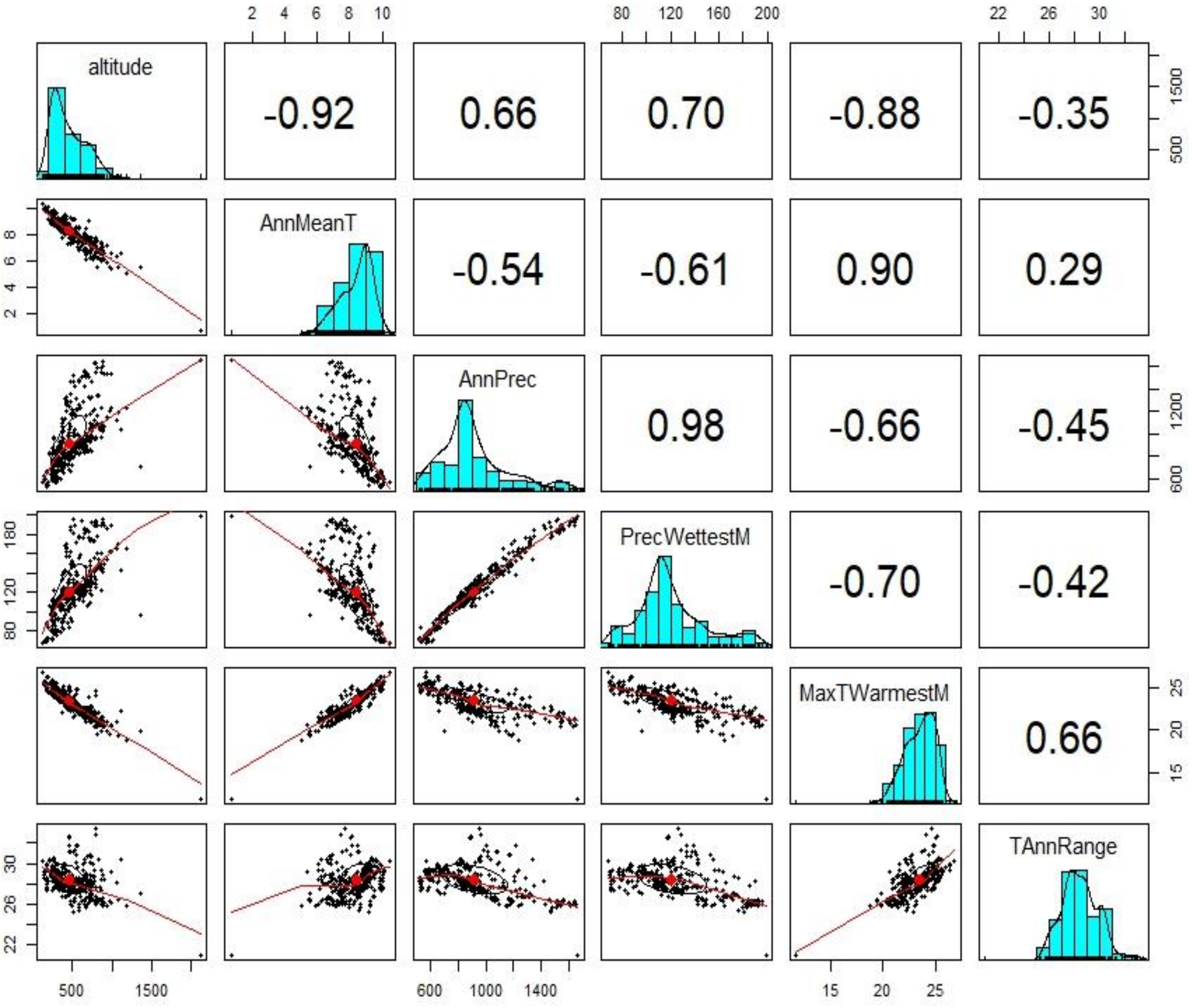


Fig. S8 Pearson´s correlations ($r^2$ values reported in the upper off diagonal quadrants) between pairs of quantitative environmental predictors, corresponding scatterplots (lower off diagonal) and distribution of single predictors as histograms with density line (diagonal).

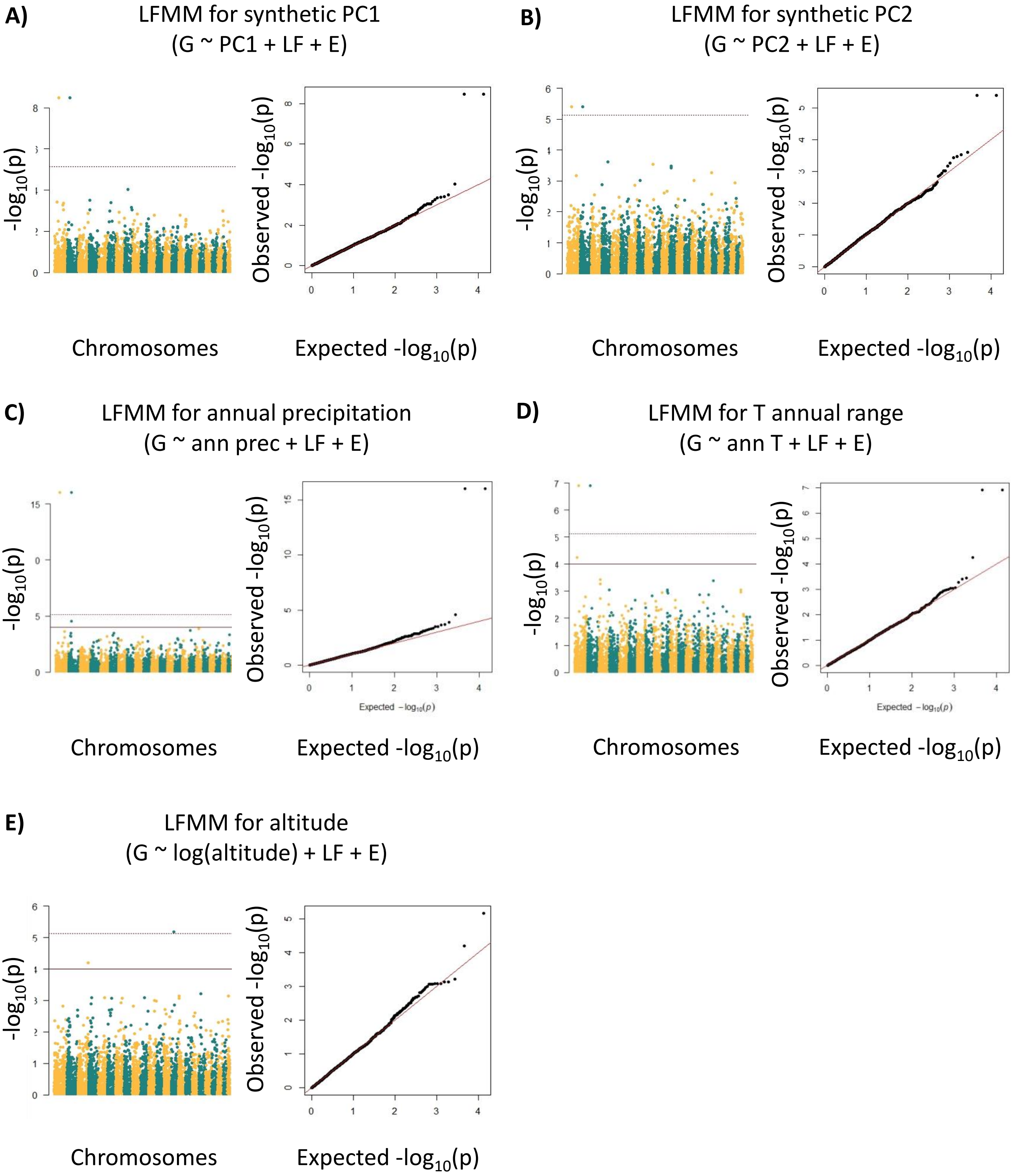


Fig. S9 Manhattan and QQ plots of LFMM. Models used are reported in the title of each pair of plots. The solid and dashed lines in Manhattan plotsrepresent a suggestive line defined at $p < 0.0001$ and a Bonferroni corrected $p < 0.05$, respectively. G = genotype matrix; LF = latent factors, PCs = first three principal components of environmental predictors.

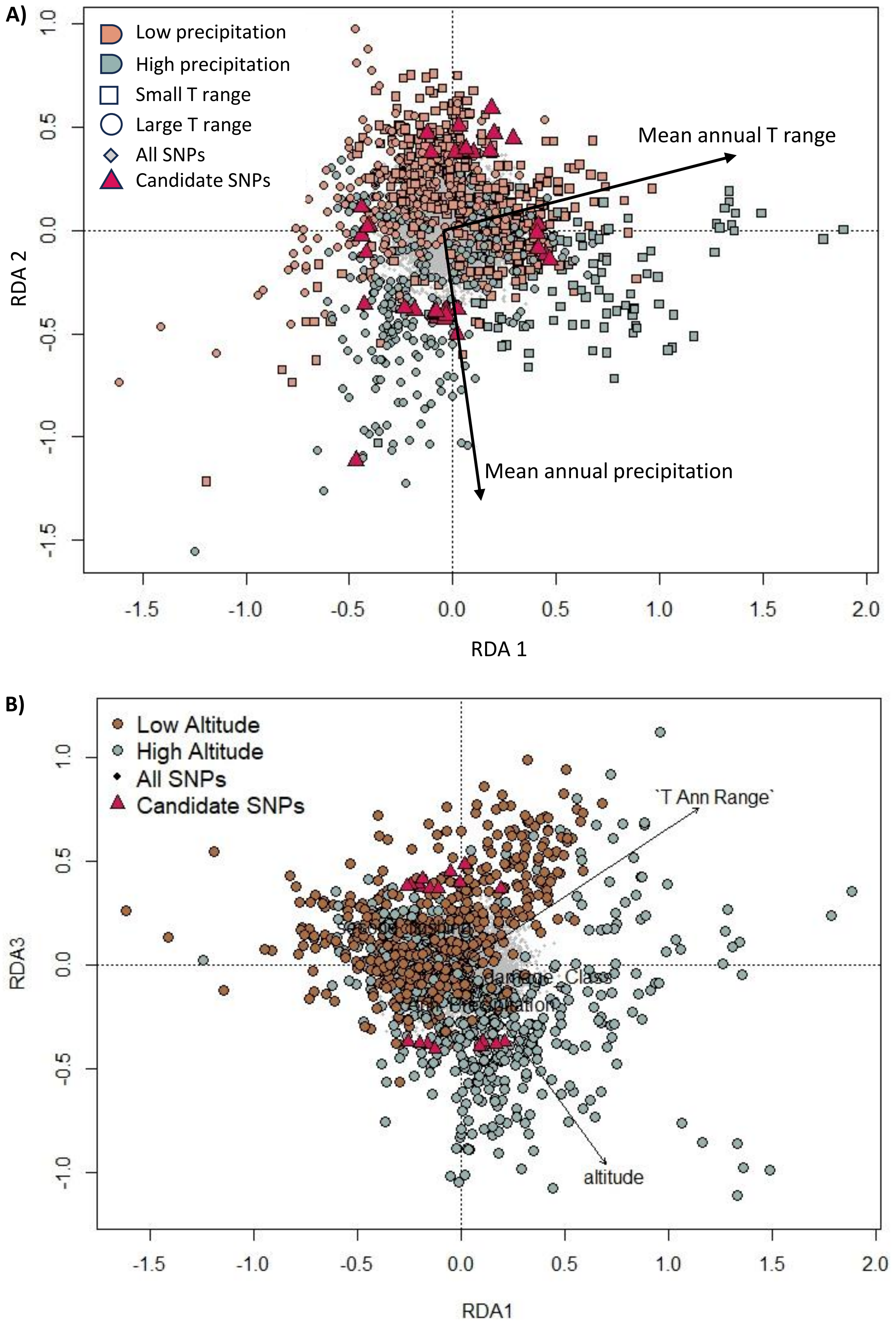

A)
Low precipitation
High precipitation
Small T range
Large T range
All SNPs
Candidate SNPs
Mean annual T range
Mean annual precipitation
RDA 2
RDA 1
B)
Low Altitude
High Altitude
All SNPs
Candidate SNPs
`T Ann Range`
altitude
RDA3
RDA1

Fig. S10 RDA plot showing the annual mean temperature and the annual mean precipitation along the first and second axes (A), and the altitude along the third axis (B). Candidate SNPs found along the axes are depicted as purple triangles.
